# Calcium-dependent protein kinases participate in RBOH-mediated sustained ROS burst during plant immune cell death

**DOI:** 10.1101/2025.09.01.672762

**Authors:** Yuta Hino, Miki Yoshioka, Hiroaki Adachi, Hirofumi Yoshioka

**Author notes:** Correspondence: Hirofumi Yoshioka.

## Abstract

The sensing of pathogen effector by an intracellular receptor called nucleotide-binding leucine-rich repeat receptor (NLR), induces a robust immune response, effector-triggered immunity (ETI). Sustained reactive oxygen species (ROS) production is accomplished by *Nicotiana benthamiana* RBOHB, an NADPH oxidase. However, molecular mechanisms connecting effector recognition and ROS production are unclear. Here, we show that calcium-dependent protein kinases (CDPKs) contribute to sustained ROS production downstream of NLR activation. We found that NbCDPK4 and NbCDPK5 directly phosphorylate NbRBOHB Ser-123 and provokes ROS production. In addition, constitutively active NbCDPKs upregulated *NbRBOHB* transcription. The phosphorylation of Ser-123 was significantly increased in a Ca^2+^-dependent manner during the ETI-like responses, which execute hypersensitive cell death. Moreover, transient expression of an autoactive helper NLR, NRC4, induced phosphorylation of Ser-123 dependent on its N-terminal conserved motif required for Ca^2+^ channel activity. These findings uncover a critical role for the NbCDPK-NbRBOHB module in regulating sustained ROS production during ETI.

## Introduction

Plant cells initiate the innate immune system following pathogen recognition. Pathogen-associated molecular pattern (PAMP) recognition by cognate cell-surface receptors induces pattern-triggered immunity (PTI), and secreted effector recognition by intracellular receptors, nucleotide-binding leucine-rich repeat receptors (NLRs), induces effector-triggered immunity (ETI) (*1*). PTI is a transient defense response targeting a broad spectrum of pathogens, while ETI is a sustained and robust response, including hypersensitive cell death response (HR cell death), against specific pathogens (*2*). Recent studies showed that there is a crosstalk between PTI and ETI. PTI signaling components are required for the initiation of ETI responses (*3*). Effector recognition reinforces PTI signaling components by transcriptional reprogramming, leading to sustained and robust defense responses (*4*). Thus, PTI and ETI share a large part of signaling pathways and second messengers, such as mitogen-activated protein kinase (MAPK) activation, Ca^2+^ influx into the cytosol, and reactive oxygen species (ROS) production (*5*).

NLRs govern the effector recognition and the induction of immune responses (*6*). NLRs are classified into three groups according to their variable domains in N-termini, including coiled-coil (CC) domain-containing NLRs (CNLs), toll/interleukin-1 receptor (TIR) domain-containing NLRs (TNLs), and resistance to powdery mildew 8 (RPW8) - like CC domain-containing NLRs (RNLs) (*7*). Activated HOPZ-ACTIVATED RESISTANCE 1 (ZAR1), an Arabidopsis CNL, forms a pentameric higher-order complex, called resistosome, which functions as a Ca^2+^ channel on the plasma membrane (*8*). Although ZAR1 is defined as a singleton NLR for both effector recognition and immune signaling, many NLRs are functionally specialized into sensor NLRs or helper NLRs (*9*). The sensor and helper NLRs work in immune receptor network architectures for adapting the diversification of pathogens. In Solanaceae plants, NLR required for cell death (NRC) subfamily proteins, known as helper NLRs, induce defense responses at the downstream of sensor NLRs (*10*). For instance, Rpi-blb2, a sensor NLR cloned from *Solanum bulbocastanum*, requires NRC4 to execute HR cell death (*10*). Activated NRC4 forms a hexameric resistosome and induce Ca^2+^ influx and HR cell death (*11*). Structural analysis reveals that the exposed α1-helix in the ZAR1 resistosome forms a funnel-like structure with a hydrophobic outer surface and a negatively charged inner surface, acting as a Ca^2+^ channel (*8, 12*). MADA motif corresponding to the ZAR1 α1-helix is conserved in helper NRCs and many CNLs (*13*). Mutation within the MADA motif of NRC4 impairs Ca^2+^ influx and HR cell death, indicating MADA motif play pivotal roles in the induction of immune responses (*11, 13*). Together, NLRs encode Ca^2+^ signals to induce downstream immune responses dependent on the N-terminal evolutionary conserved motif.

ROS are required for development and stress responses in plants (*14*). Hydrogen peroxide acts as a second messenger through oxidative post-translational modification of cysteine residues (*15*). ROS production is strictly regulated, because excess ROS cause oxidative stress for plant cells. Upon pathogen attack, the plasma membrane-localized NADPH oxidase, respiratory burst oxidase homolog (RBOH), contributes to ROS production in the apoplastic region (*16*). In *Nicotiana benthamiana*, NbRBOHB is responsible for plant immune ROS production, which is required for the induction of HR cell death and defense against *Phytophthora infestans* (*16*). RBOH is a six-transmembrane protein, and its activity is regulated by post-translational modification of the N- and C-terminal regions exposed to cytosol (*17*). For instance, phosphorylation in the N-terminal region of AtRBOHD by receptor-like cytoplasmic kinase (RLCK) activates RBOH upon PAMP recognition (*18*). A transmembrane kinase, cysteine-rich receptor kinase 2 (CRK2), also phosphorylates the C-terminal region of RBOH and plays a positive role in flg22-induced ROS production (*19*). Furthermore, phosphatidic acid (PA) binding to AtRBOHD positively regulates the RBOH-mediated ROS production by enhancing their protein accumulation (*20, 21*).

RBOH activity is regulated in a Ca^2+^-dependent manner. Ca^2+^-chelator and Ca^2+^ channel blocker inhibit RBOH-dependent ROS production (*22*). Interaction between Ca^2+^ and EF-hand motifs in the N-terminus of RBOH are required for inducing ROS production (*23*). Calcium-dependent protein kinases (CDPKs) are serine/threonine protein kinases containing a Ca^2+^-binding calmodulin-like domain (*23*). CDPKs regulate plant development, pollen tube growth, stomatal movements, and stress responses (*24*). CDPKs directly phosphorylate the N-termini of RBOH and activate ROS production. In potato tuber, StCDPK4 and StCDPK5 phosphorylate StRBOHB Ser-82 and Ser-97, and induce ROS production (*25*). In Arabidopsis, AtCPK4, AtCPK5, AtCPK6, and AtCPK11 are responsible for the phosphorylation of AtRBOHD (*18, 26*). Transgenic potato plants containing constitutively active StCDPK5 under the control of a pathogen-inducible promoter show enhanced ROS accumulation and resistance against *P. infestans*, supporting the importance of CDPK-mediated ROS production in defense against near-obligate biotrophic pathogens (*27*).

In addition to post-translational modification of RBOH, transcriptional reprogramming of RBOH through the MAPK-WRKY pathway is required for the sustained ROS production during ETI (*28*). In contrast, overexpression of RBOH via Agrobacterium did not induce ROS production (*27*), supporting that sustained ROS production during ETI is accomplished by both transcriptional and post-translational regulation of RBOH. Given that cytosolic Ca^2+^ elevation occurred after NLR activation, CDPK might play a pivotal role in regulating downstream immune responses such as ROS production. However, the molecular mechanisms connecting NLR activation to defense responses have been still unknown.

In this study, we show that NbCDPK4 and NbCDPK5, classified into subgroup I CDPK, directly phosphorylated NbRBOHB Ser-123 and regulated sustained ROS production during ETI in *N. benthamiana*. The phosphorylation of Ser-123 was significantly induced during an ETI-like response accompanied by HR cell death, but not during PTI. Moreover, transient expression of an autoactive helper NLR induced NbRBOHB-dependent ROS production and Ser-123 phosphorylation in Ca^2+^-dependent manner. Our results pave the way to reveal the regulatory molecular mechanisms connecting the helper NLR activation and downstream immune responses.

## RESULTS

### NbCDPK4 and NbCDPK5 interact with NbRBOHB and generate ROS production

Multiple CDPKs participate in the post-translational activation of RBOH (*18, 25*). To estimate CDPKs that are involved in the regulation of NbRBOHB, we conducted a phylogenetic analysis of *N. benthamiana* CDPKs with selected plant CDPKs reported in a previous study (*29*). This analysis identified NbCDPK4 (Niben101Scf09867g01003.1), NbCDPK5 (Niben101Scf01191g06012.1), which are the closest homologs of potato StCDPK4 and StCDPK5 respectively, and NbCDPK9 (Niben101Scf07616g00008.1), NbCDPK10 (Niben101Scf04216g07017.1), NbCDPK11 (Niben101Scf05534g01007.1), which are classified into the same clade with AtCPK4 and AtCPK11 (Fig. 1A). The CDPK family protein generally consists of an N-terminal variable (V) domain, a protein kinase (K) domain, a junction (J) domain, and a calmodulin-like (C) domain including four EF-hand motifs (Fig. 1B) (*30*). NbCDPK4, NbCDPK5, NbCDPK9, NbCDPK10, and NbCDPK11 encode conventional CDPK containing VKJC domains. NbCDPK4 and NbCDPK5 carry estimated myristoylation and S-palmitoylation (S-acylation) sites (Gly-2/Cys-5) at their N termini, which are required for plasma membrane anchoring, but NbCDPK9, NbCDPK10, and NbCDPK11 do not (fig S1).

**Fig. 1.**
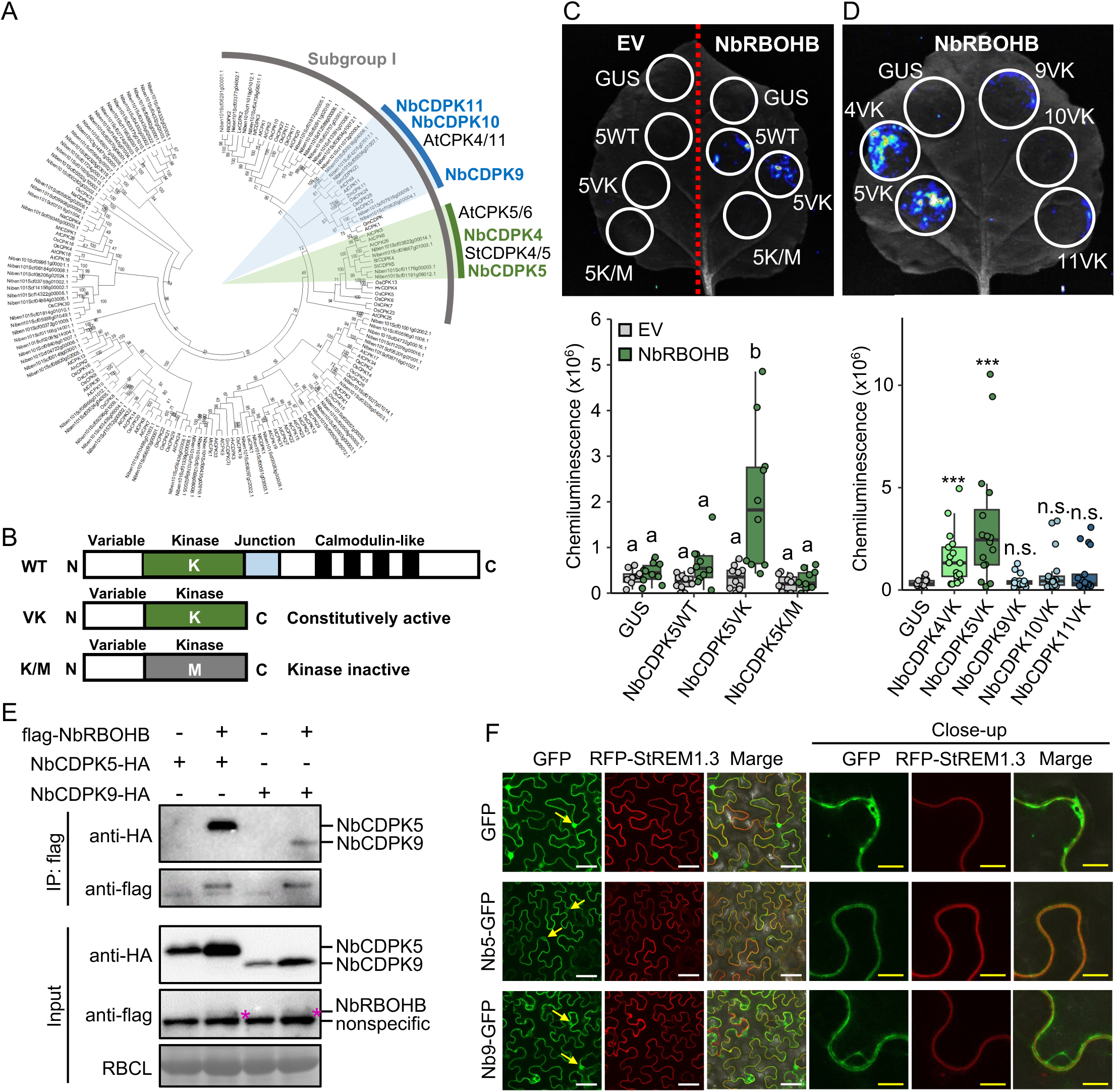
NbCDPK4 and NbCDPK5 interact with NbRBOHB and generate ROS production. (A) Phylogenetic tree of *Nicotiana benthamiana* NbCDPKs with selected plant CDPKs; Arabidopsis (At), rice (Os), soybean (Gm), potato (St), barley (Hv), tobacco (Nt), coyote tobacco (Na), tomato (Le) and grapevine (ACPK1), maize (Zm), alfalfa (Mt), ice plant (Mc) and peanut (Ah) (*29*). The clade including NbCDPK4 and NbCDPK5 is indicated by the green area, and the clade including NbCDPK9, NbCDPK10 and NbCDPK11 is indicated by the blue area. Predicted NbCDPK sequences were obtained from iTAK Plant Transcription factor & Protein Kinase Identifier and Classifier. Phylogenetic analyses using the neighbor-joining method were done in MEGA7. (B) Schematic structures of NbCDPK with variable, kinase, junction, and calmodulin-like domains. K/M is a truncated kinase inactive variant with substitution in Lys (K) of the ATP binding motif to Met (M). (C) ROS production in *N. benthamiana* leaves expressing NbCDPK5 variants. *A. tumefaciens* harboring NbCDPK5 variants or GUS, and NbRBOHB or empty vector (EV) were mixed 1:4 (OD600 = 0.5) and infiltrated. To detect ROS generation, leaves were infiltrated with 0.5 mM L-012 solution two days after agroinfiltration. Letters represent each significance group, determined through Tukey’s multiple range test. (D) ROS production induced by expressing constitutively active variants (VK) of subgroup I NbCDPKs with NbRBOHB. *A. tumefaciens* harboring NbCDPK and NbRBOHB were mixed 1:4, and were infiltrated. Asterisks indicate statistically significant differences compared to GUS control (t test, ***P < 0.005). (E) Interaction between NbCDPKs and NbRBOHB. Co-immunoprecipitation was performed with extracts from *N. benthamiana* leaves co-expressing HA-tagged NbCDPK5 or NbCDPK9 with FLAG-tagged NbRBOHB. µMACS magnetic beads with anti-FLAG antibody were used for immunoprecipitation, and anti-HA and anti-FLAG antibodies were used to detect the related proteins in the immunoprecipitants. Magenta pentagrams indicate the signals of NbRBOHB. Protein loads were monitored by Ponceau 4R staining of the bands corresponding to Rubisco large subunit (RBCL). (F) Subcellular localization of NbCDPK5 and NbCDPK9. GFP-tagged NbCDPK5 or NbCDPK9 were transiently co-expressed with RFP-StREM1.3, and observed under the confocal microscope 3 days after agroinfiltration. Arrows indicate the nucleus. White bars = 50 µm, yellow bars = 10 µm.

To investigate whether NbCDPK5 has a capacity to activate ROS production, we constructed variants of NbCDPK5. VK mutant is a truncated variant lacking J and C domains and is used as a constitutively active kinase mutant (Fig. 1B) (*31, 32*). K/M is a VK variant with a Lys-to-Met mutation at the ATP binding site (K110), disrupting its protein kinase activity (Fig. 1B, fig. S1) (*33, 34*). We measured ROS production after expressing NbCDPK5 variants (wild-type: WT, VK, and K/M) and b-glucuronidase (GUS) protein (negative control) in *N. benthamiana* leaves (Fig. 1C). NbCDPK5VK, but not NbCDPK5WT, NbCDPK5K/M and GUS, significantly induced ROS production when they are co-expressed with NbRBOHB (Fig. 1C). We further generated constitutively active VK mutants of NbCDPK4, NbCDPK9, NbCDPK10, and NbCDPK11. NbCDPK4VK and NbCDPK5VK induced ROS accumulation when expressed with NbRBOHB, while NbCDPK9VK, NbCDPK10VK, and NbCDPK11VK did not (Fig. 1D). These results indicate that NbCDPK4 and NbCDPK5 potentially activate NbRBOHB-dependent ROS production in *N. benthamiana*.

CDPK physically interacts with RBOH on the plasma membrane (*35*). To examine the interaction between NbCDPKs and NbRBOHB, we performed a co-immunoprecipitation (co-IP) assay. In this assay, we used NbCDPK5 and NbCDPK9 as representative CDPKs with or without myristylation and S-acylation sites, respectively. This co-IP experiment revealed that both of NbCDPK5 and NbCDPK9 interact with NbRBOHB. Notably, NbCDPK5-NbRBOHB interaction was stronger than that of NbCDPK9 (Fig. 1E). Myristylation- and S-acylation-mediated plasma membrane localization of CDPK is required for the interaction between CDPK and RBOH (*35*). To confirm the subcellular localization of NbCDPKs, the fluorescence from GFP-fused NbCDPKs were observed under confocal microscopy (Fig. 1F, fig. S2). In this experiment, we used RFP-StREM1.3 as a plasma membrane marker (*36*). NbCDPK4-GFP and NbCDPK5-GFP were mainly localized to the plasma membrane, while NbCDPK9-GFP, NbCDPK10-GFP, and NbCDPK11-GFP were localized in the cytosol (Fig. 1F, fig. S2). GFP-fused NbCDPK5, NbCDPK9, NbCDPK10, and NbCDPK11 were also localized in the nucleus. In addition, NbCDPK4 and NbCDPK5 were highly accumulated in the microsome fractions, but NbCDPK9, NbCDPK10 and NbCDPK11 mainly existed in the soluble fractions (fig. S3). These results indicate that NbCDPK4 and NbCDPK5 are mainly localized to the plasma membrane, and presumably interact with NbRBOHB on the plasma membrane.

### NbCDPK4 and NbCDPK5 activate NbRBOHB by direct phosphorylation and transcriptional reprogramming

In *S. tuberosum*, StCDPK4 and StCDPK5 activate ROS production by phosphorylating Ser-82 and Ser-97 of StRBOHB (*25*). Phosphorylation motifs at Ser-82 and Ser-97 of StRBOHB are conserved in NbRBOHB at Ser-123 and Ser-138 (Fig. 2A). To determine whether Ser-123 in NbRBOHB is phosphorylated by NbCDPKs, we prepared an anti-phosphopeptide antibody against peptides including phospho-Ser-123 (pS123). Because the anti-StRBOHB pS97 antibody gave non-specific signals in a previous study and amino acid sequence including NbRBOHB Ser-138 showed high similarity, we could not prepare a specific antibody against peptides including pS138 (*25*). We purified NbCDPK5 protein expressed in *Escherichia coli.* The recombinant NbCDPK5 protein phosphorylated Ser-123 site in the N-terminal fragment of NbRBOHB in a Ca^2+^-dependent manner *in vitro* (Fig. 2B). Next, we examined whether NbCDPK5 phosphorylates the Ser-123 of NbRBOHB in plants. NbCDPK5VK phosphorylated NbRBOHB Ser-123 compared with GUS, NbCDPK5WT, and NbCDPK5K/M, when expressed in *N. benthamiana* leaves (Fig. 2C). Consistent with ROS accumulation level, NbCDPK4VK and NbCDPK5VK strongly phosphorylated NbRBOHB Ser-123, while NbCDPK9VK, NbCDPK10VK and NbCDPK11VK did not (Fig. 1D, Fig. 2D). To validate the degree to which the Ser-123 phosphorylation contributes to NbRBOHB-mediated ROS production, we generated an alanine-substituted mutant of NbRBOHB (NbRBOHB^S123A^). ROS production caused by NbCDPK5VK expressed with NbRBOHB^S123A^ was much less than that with NbRBOHB^WT^ (Fig. 2E). These results indicate that NbCDPK5 activates NbRBOHB-dependent ROS production by direct phosphorylation of NbRBOHB Ser-123.

**Fig. 2.**
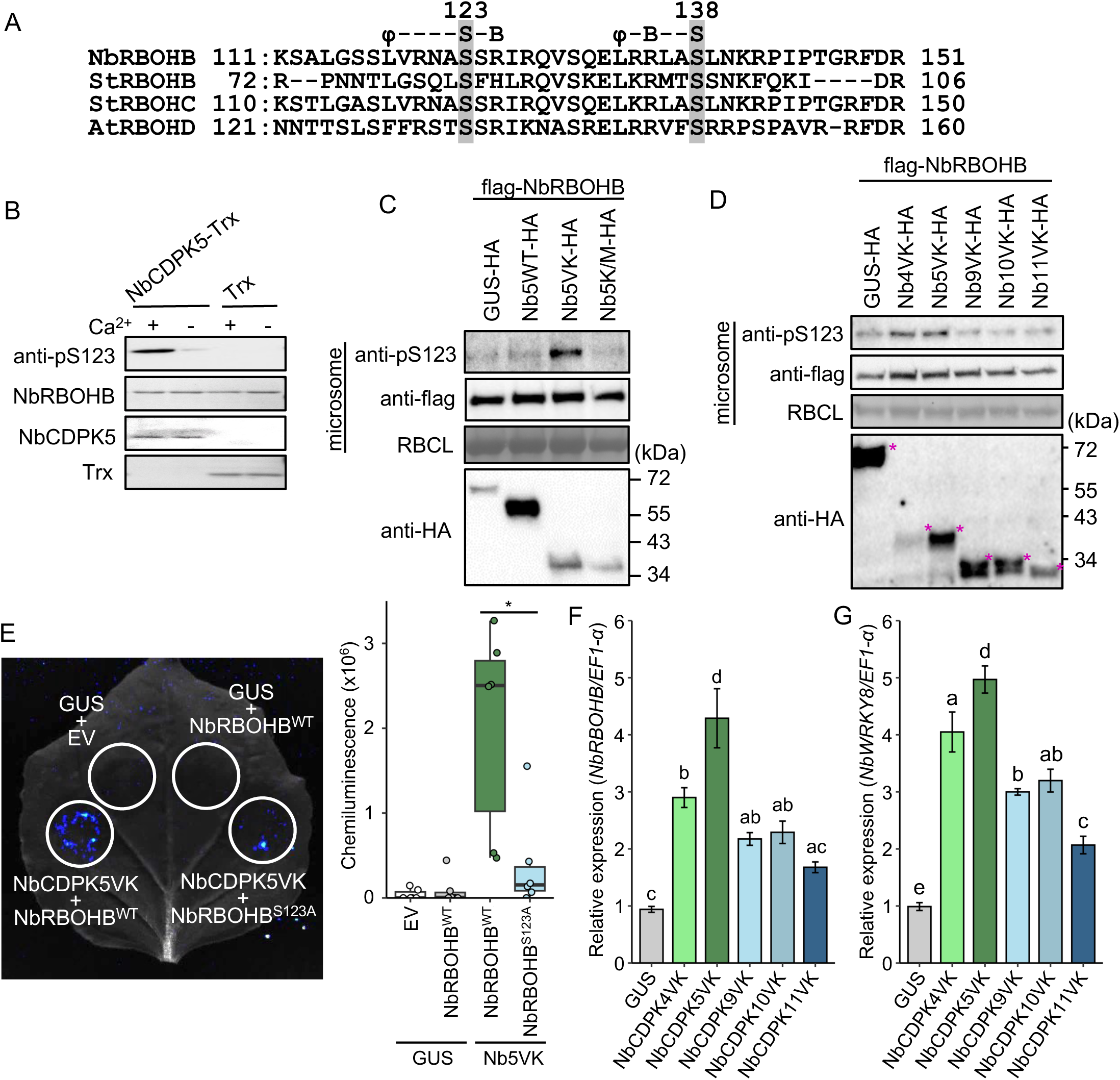
NbCDPK4 and NbCDPK5 activate NbRBOHB by direct phosphorylation and transcriptional reprogramming.(A) Conservation of potato StCDPK5-mediated StRBOHB phosphorylation sites (Ser-82 and Ser-97 corresponding to Ser-123 and Ser-138 of *Nicotiana benthamiana* NbRBOHB). The amino acid sequences of plant immune RBOHs, Arabidopsis AtRBOHD, StRBOHB, StRBOHC and NbRBOHB were aligned by ClustalW. A consensus motif for CDPK-mediated phosphorylation is indicated upper the sequences of RBOHs. (B) Ca^2+^-dependent phosphorylation of NbRBOHB Ser-123 by NbCDPK5 *in vitro*. Purified N-terminal peptides of NbRBOHB were used as substrates for Trx-NbCDPK5. Phosphorylation of NRBOHB was detected using anti-pS123 antibody. Protein loads were monitored by MemCode staining. (**C** and **D**) Phosphorylation of NbRBOHB Ser-123 by constitutively active NbCDPK5 (C) and subgroup I NbCDPKs (D) in plants. NbCDPK variants or GUS were co-expressed with NbRBOHB (mixed 1:4). Microsomal fractions were prepared two days after agroinfiltration. Immunoblot analyses were performed using anti-pS123, anti-flag and anti-HA antibodies. Protein loads were monitored by MemCode staining of the bands corresponding to Rubisco large subunit (RBCL). Magenta pentagrams indicate the signals of anti-HA antibody. (**E**) Effects of mutations of Ser-123 on the ROS production induced by NbCDPK5VK. *A. tumefaciens* harboring NbCDPK5 and NbRBOHB variants were mixed 1:4, and were infiltrated. To detect ROS generation, leaves were infiltrated with 0.5 mM L-012 solution two days after agroinfiltration. Asterisks indicate statistically significant differences (t test, *P < 0.05). (**F** and **G**) Expression of *NbRBOHB* (F) and *NbWRKY8* (G) induced by constitutively active variants (VK) of subgroup I NbCDPKs. Total RNAs were extracted from leaves two days after agroinfiltration, and were used for RT-qPCR. Letters represent each significance group, determined through Tukey’s multiple range test.

Because these NbCDPKs were localized in nucleus, we hypothesized that NbCDPKs participate in the transcriptional reprogramming of *NbRBOHB*. To test this hypothesis, transcription levels of *NbRBOHB* were validated by RT-qPCR. VK mutants of NbCDPK4, NbCDPK5, NbCDPK9, NbCDPK10, and NbCDPK11 up-regulated *NbRBOHB* expression (Fig. 2F), indicating that NbCDPKs support ROS burst by transcriptional reprogramming of *NbRBOHB*. In addition, all tested VK mutants also induced *NbWRKY8* expression, which is a transcription factor regulating promoter activity of *NbRBOHB* (Fig. 2G). These results suggest that NbCDPKs might induce *NbRBOHB* expression through the transcriptional regulation of *NbWRKY8*.

### The phosphorylation of NbRBOHB Ser-123 was strongly induced during immune responses associated with cell death

To investigate whether the phosphorylation of NbRBOHB Ser-123 is induced during plant immune responses, we investigated the phosphorylation of Ser-123 in response to PAMP treatment or effector expression. Estradiol-induced expression of *AVRblb2* in Rpi-blb2 transgenic *N. benthamiana* leaves induced the phosphorylation of NbRBOHB Ser-123 (Fig. 3A). NbRBOHB Ser-123 phosphorylation was also induced in response to INF1 at 6, 12, and 24 hours after the treatment (Fig. 3B, fig. S4). INF1 treatment induces two-peak ROS production, transient first peak (around an hour after treatment) and transcriptional reprogramming-dependent continuous second peak (*28, 37*). At early time points (10, 30 and 60 minutes after the INF1 treatment), the phosphorylation of Ser-123 was not detected (Fig. 3B). In addition, flg22, which induces single-peak transient ROS production in *N. benthamiana*, did not induce the phosphorylation of NbRBOHB Ser-123, even though Ser-337, a predicted phosphorylation site by RLCK (*18, 38*), was phosphorylated (Fig. 3C). These results indicate that the phosphorylation of NbRBOHB Ser-123 was dominantly induced during ETI-like robust immune responses.

**Fig. 3.**
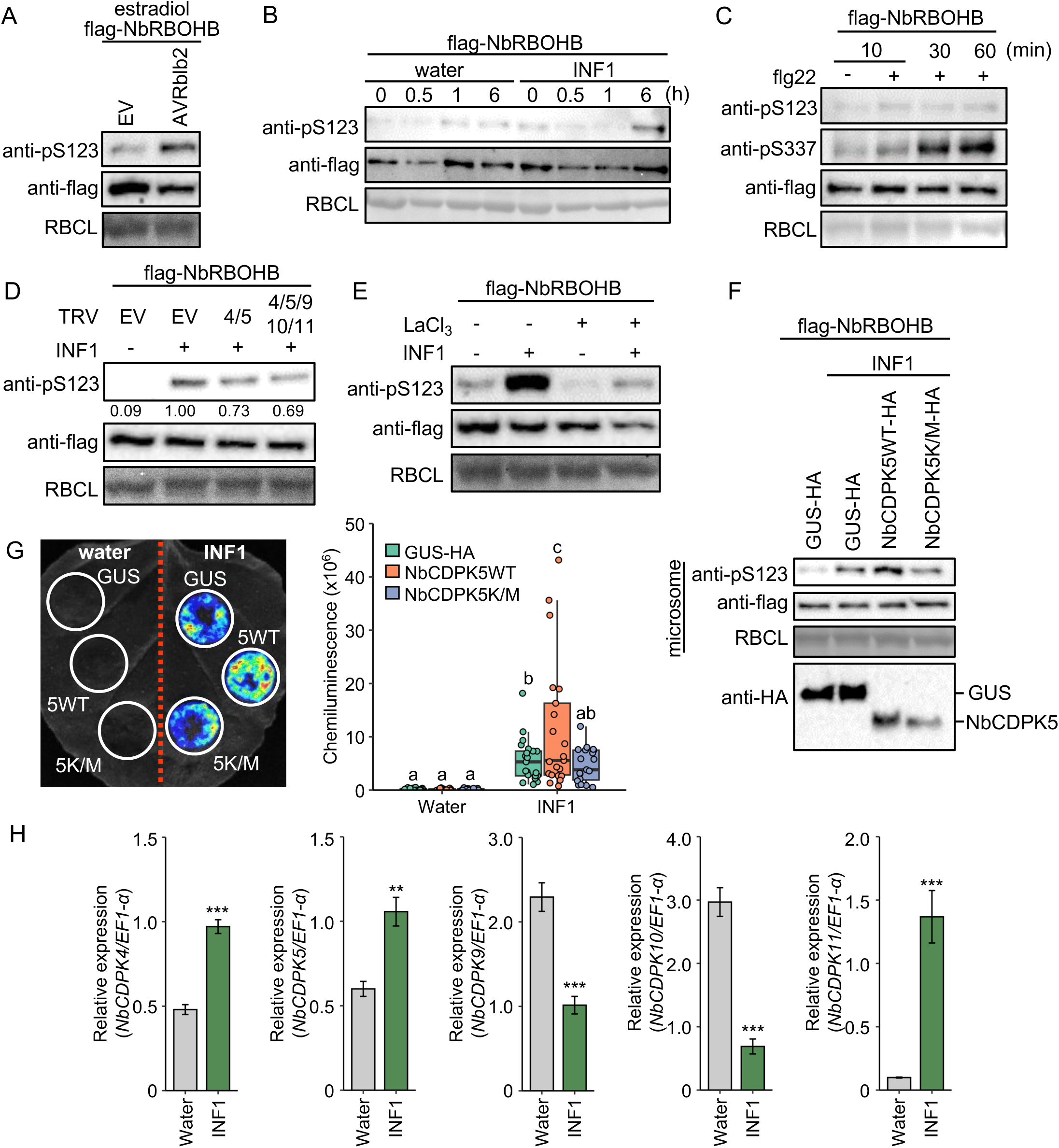
The phosphorylation of NbRBOHB Ser-123 was strongly induced during ETI. (**A** to **C**) Phosphorylation of NbRBOHB Ser-123 in response to (A) AVRblb2, (B) INF1 peptide, and (C) flg22 peptide. NbRBOHB was transiently expressed by agroinfiltration. Two days after agroinfiltration, the leaves were treated with elicitors. Microsomal fractions were prepared at an indicated time after treatment. Immunoblot analyses were performed using anti-pS123, anti-pS337 and anti-flag antibodies. Protein loads were monitored by MemCode staining of the bands corresponding to RBCL. (D) Ca^2+^-channel blocker impaired the phosphorylation of NbRBOHB Ser-123. NbRBOHB-expressing leaves were infiltrated with 5 mM LaCl3 an hour before INF1 treatment. Microsomal fractions were prepared 6 hours after INF1 treatment. (E) Silencing of *NbCDPKs* attenuated the phosphorylation of NbRBOHB Ser-123. Microsomal fractions were prepared from NbCDPKs multiple silenced *N. benthamiana* 6 hours after INF1 treatment. The relative intensity of anti-pS123 was indicated below the top panel. (F) Overexpression of NbCDPK5 enhanced the phosphorylation of NbRBOHB Ser-123. NbCDPK5WT, NbCDPK5K/M, or GUS were co-expressed with NbRBOHB by agroinfiltration (mixed 1:4) for two days. Microsomal fractions were prepared 6 hours after INF1 treatment. (G) Effects of NbCDPK5 overexpression on the ROS production induced by INF1. *A. tumefaciens* harboring NbCDPK5 variants were infiltrated. Leaves were treated with INF1 two days after agroinfiltration. To detect ROS generation, leaves were infiltrated with 0.5 mM L-012 solution 6 hours after INF1 treatment. Letters represent each significance group, determined through Tukey’s multiple range test. (H) Expression of *NbCDPKs* in response to INF1. Total RNAs were extracted from leaves 24 hours after INF1 treatment and were used for RT-qPCR. Asterisks indicate statistically significant differences compared to water treatment (t test, **P < 0.01, ***P < 0.005).

Next, we examined whether NbCDPKs play an important role in the phosphorylation of NbRBOHB Ser-123 in response to the INF1 infiltration. The phosphorylation of NbRBOHB Ser-123 was attenuated by about 30% in both *NbCDPK4/5*- and *NbCDPK4/5/9/10/11*-silenced plants (Fig. 3D). Moreover, INF1-induced phosphorylation of NbRBOHB Ser-123 was inhibited in the presence of LaCl3, a Ca^2+^-channel blocker (Fig. 3E). These results suggest that NbCDPK4, NbCDPK5, and other kinases participate in the phosphorylation of NbRBOHB Ser-123, and at least this phosphorylation requires Ca^2+^ influx.

To investigate whether NbCDPK5 is involved in immune responses induced by INF1, we examined ROS accumulation and the phosphorylation of NbRBOHB under NbCDPK5 overexpressing conditions. NbCDPK5WT, but not NbCDPK5K/M, overexpressing plants showed enhanced phosphorylation of NbRBOHB Ser-123 in response to INF1 (Fig. 3F). Consistent with the phosphorylation levels, INF1-induced ROS accumulation was increased by NbCDPK5WT overexpression (Fig. 3G). Moreover, transcription of *NbCDPK5* was induced in response to INF1 (Fig. 3H), suggesting transcriptional increase in the NbCDPK5 contributes to the robust ROS production during ETI-like response. In addition, expression levels of *NbCDPK4* and *NbCDPK11* also increased by INF1, while those of *NbCDPK9* and *NbCDPK10* rather decreased (Fig. 3H). In contrast to INF1-induced ROS production, flg22-induced ROS production was not affected by NbCDPK5WT overexpression (fig. S5), which was not contradict with Ser-123 phosphorylation status by flg22 (Fig 3C). These results indicate that NbCDPK5 positively regulates sustained ROS burst in response to INF1, but not the transient ROS burst by flg22.

### The phosphorylation of NbRBOHB Ser-123 was induced during spontaneous cell death

To further investigate the relationship between the phosphorylation of NbRBOHB and cell death, we examined the phosphorylation of NbRBOHB at Ser-123 and Ser-337 in response to several signal inputs that induce spontaneous cell death. For instance, transient expression of MEK2^DD^, a constitutively active mutant of immune-related MAPKK, induces NbRBOHB-dependent ROS production and cell death (*28*). NbRBOHB was phosphorylated at both Ser-123 and Ser-337 at 24 and 36 hours after agroinfiltration to express MEK2^DD^ (Fig. 4A). Bax, inducing mitochondrial dysfunction-mediated apoptosis (*39*), also induced Ser-123 phosphorylation (Fig. 4B). On the other hand, chemical-induced cell death caused by ethanol did not induce Ser-123 phosphorylation (Fig. 4C). These results indicate that the phosphorylation of NbRBOHB at Ser-123 is induced during biological spontaneous cell death, especially regulated by MAPK cascade and mitochondrial dysfunction.

**Fig. 4.**
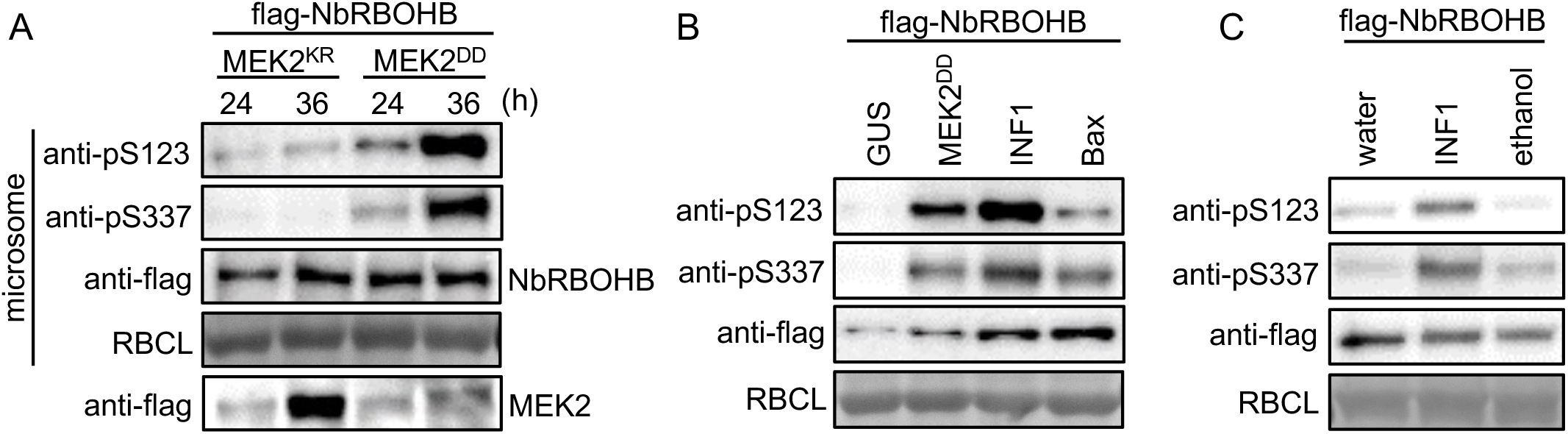
The phosphorylation of NbRBOHB Ser-123 was induced during spontaneous cell death. (A) Phosphorylation of NbRBOHB Ser-123 induced by constitutively active MAPKK. *A. tumefaciens* harboring NtMEK2 and NbRBOHB were mixed 1:1, and were infiltrated. Microsomal fractions were prepared at an indicated time after agroinfiltration. Immunoblot analyses were performed using anti-pS123, anti-pS337 and anti-flag antibodies. Protein loads were monitored by MemCode staining of the bands corresponding to RBCL. (B) Phosphorylation of NbRBOHB Ser-123 induced by Bax. NtMEK2^DD^, INF1, Bax or GUS were co-expressed with NbRBOHB by agroinfiltration (mixed 1:1). Microsomal fractions were prepared a day after agroinfiltration. Protein loads were monitored by Ponceau 4R staining of the bands corresponding to RBCL. (C) Ethanol did not induce phosphorylation of NbRBOHB Ser-123. NbRBOHB was transiently expressed via Agrobacterium. Two days after agroinfiltration, the leaves were treated with 20% (v/v) ethanol or INF1. Microsomal fractions were prepared six hours after treatment. Protein loads were monitored by Ponceau 4R staining of the bands corresponding to RBCL.

### CDPKs activate NbRBOHB downstream of helper NLR

Upon effector recognition, resistosome formation of helper NLR and its Ca^2+^-channel activity are required to proceed with the HR cell death (*11*). We investigated the involvement of the CDPK-RBOH module in executing HR cell death following helper NLR activation. Consistent with the previous report, transient expression of autoactive mutant NRC4^D478V^ induced HR cell death, which was impaired by L9E mutation within the MADA motif located at the N-terminus of NRC4 (*13*) (fig. S6). The NRC4^L9E/D478V^ can form the activated resistosome, but its Ca^2+^-channel activity is terminated (*11*). NRC4^D478V^ expression induced apoplastic ROS accumulation and the phosphorylation of NbRBOHB Ser-123 at 18 and 24 hours after agroinfiltration, and the L9E mutation impaired these responses (Fig. 5A, B). Moreover, LaCl3 inhibited both apoplastic ROS accumulation and phosphorylation of NbRBOHB Ser-123 triggered by NRC4^D478V^ (Fig. 5C, D). To avoid the effect of LaCl3 treatment on transient expression via *Agrobacterium tumefaciens*, LaCl3 was infiltrated 12 hours after agroinfiltration in this experiment. We confirmed that NRC4 and RBOHB proteins accumulate similarly with or without the LaCl3 treatment (Fig. 5D). These results indicate that Ca^2+^ is a key signal messenger downstream of the helper NRC4 to induce ROS accumulation and NbRBOHB phosphorylation.

**Fig. 5.**
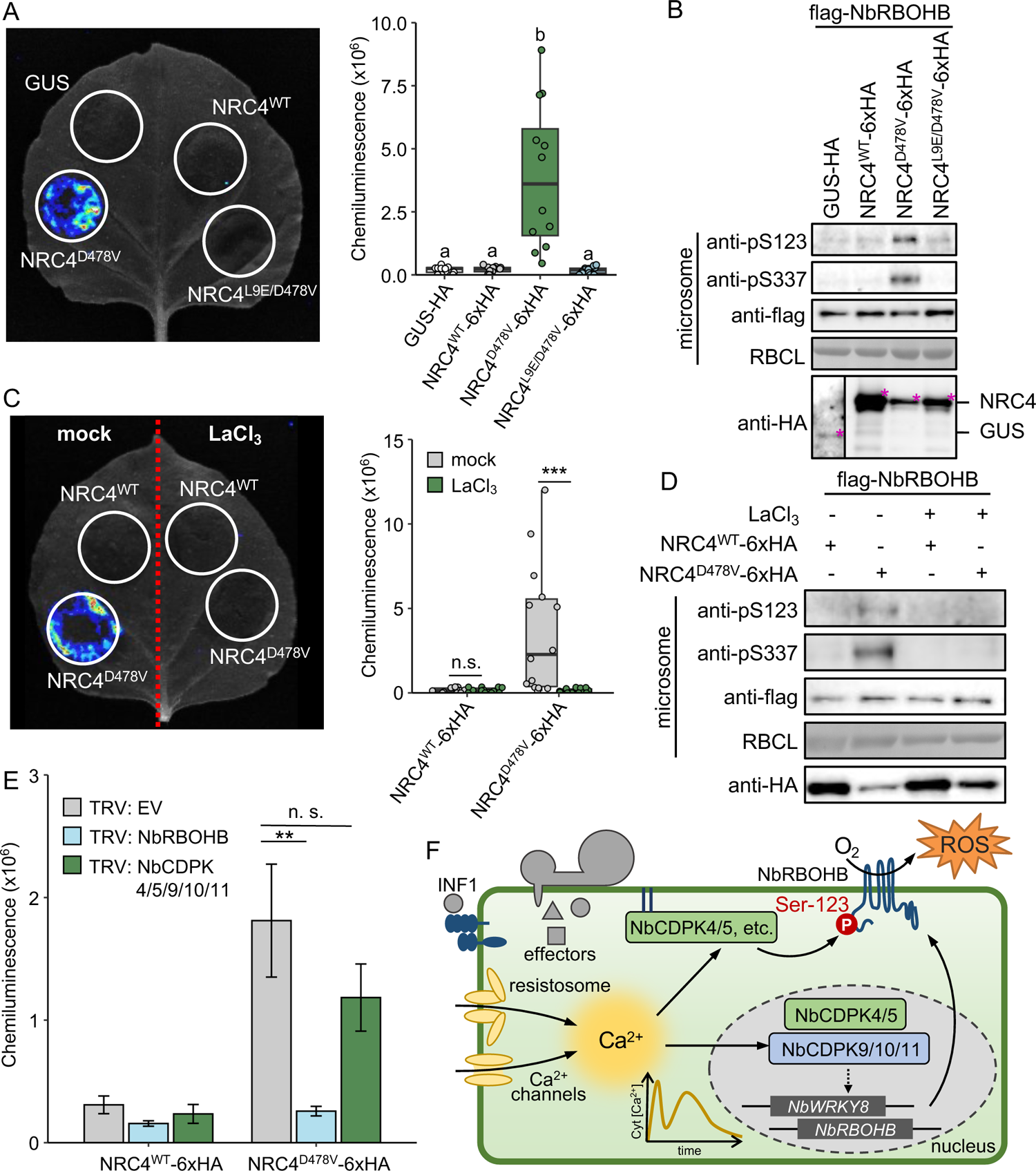
CDPKs activate NbRBOHB downstream of autoactive helper NLR. (A) ROS production induced by expression of autoactive NRC4^D478V^, but not by inactive NRC4^L9ED478V^. *A. tumefaciens* harboring NRC4 variants (OD600 = 0.2) were infiltrated. To detect ROS generation, leaves were infiltrated with 0.5 mM L-012 solution 18 hours after agroinfiltration. Letters represent each significance group, determined through Tukey’s multiple range test. (B) Phosphorylation of NbRBOHB Ser-123 after expression of NRC4 variants. NRC4 variants were co-expressed with NbRBOHB by agroinfiltration (mixed 1:4; OD600 = 0.5). Microsomal fractions were prepared 24 hours after agroinfiltration. Immunoblot analyses were performed using anti-pS123, anti-pS337, anti-flag and anti-HA antibodies. Protein loads were monitored by ponceau 4R staining of the bands corresponding to RBCL. Magenta pentagrams indicate the signals of anti-HA antibody. (C) Ca^2+^-channel blocker impaired ROS production induced by NRC4^D478V^. ROS production was measured by chemiluminescence mediated by 0.5 mM L-012 18 hours after infiltration of *A. tumefaciens*. 5 mM LaCl3 was infiltrated 12 hours after agroinfiltration. Asterisks indicate statistically significant differences compared to mock treatment (t test, ***P < 0.005). (D) Ca^2+^-channel blocker impaired phosphorylation of NbRBOHB Ser-123 induced by NRC4^D478V^. NRC4 variants were co-expressed with NbRBOHB by agroinfiltration (mixed 1:4; OD600 = 0.5). 5 mM LaCl3 was infiltrated, and microsomal fractions were prepared 12 and 24 hours after agroinfiltration, respectively. Protein loads were monitored by ponceau 4R staining of the bands corresponding to RBCL. (E) NRC4^D478V^ induced NbRBOHB-dependent ROS production. *A. tumefaciens* harboring NRC4^D478V^ or GUS (OD600 = 0.2) were infiltrated into *NbRBOHB*-silenced or *NbCDPKs*-silenced *N. benthamiana* leaves. Asterisks indicate statistically significant differences compared to TRV control (t test, **P < 0.01). (F) Model of the regulatory mechanism of sustained ROS bursts by CDPKs during ETI-like responses. Recognition of effectors induces continuous elevation of Ca^2+^ concentrations in cytosol by activating helper NLR resistosome and Ca^2+^ channels. The NbCDPK4 and NbCDPK5, etc. localizing to plasma membrane directory phosphorylate NbRBOHB Ser-123, followed by ROS production under high Ca^2+^ concentrations. NbCDPK4, NbCDPK5, NbCDPK9, NbCDPK10, and NbCDPK11 increase transcriptional levels of NbRBOHB and NbWRKY8, resulting in the supply of newly synthesized NbRBOHB.

Next, we investigated whether NbRBOHB contributes to apoplastic ROS accumulation induced by NRC4^D478V^. The NRC4^D478V^-mediated ROS accumulation was inhibited by *NbRBOHB* silencing (Fig. 5E). We found no significant difference in apoplastic ROS accumulation between *NbCDPK4/5/9/10/11*-silenced plants and TRV-control (Fig. 5E). These results indicate that NbRBOHB plays a pivotal role in apoplastic ROS accumulation mediated by the NRC4 resistosome, and the NbRBOHB activity is presumably regulated not only by the CDPKs but also by other protein kinases.

## DISCUSSION

Post-translational activation of RBOH is crucial for regulating ROS burst during plant immune responses. We previously reported that the phosphorylation of StRBOHB by StCDPK4 and StCDPK5 activates ROS burst, and MAPK-WRKY pathway-dependent transcriptional reprogramming is required for sustained ROS burst during ETI (*25, 28*). However, molecular mechanisms regulating RBOH activity to maintain sustained ROS burst upon effector recognition have been poorly understood. Here, we showed that the phosphorylation of NbRBOHB Ser-123 by NbCDPK4 and NbCDPK5 played a dominant role in the regulation of ROS production downstream of NLR activation (Fig. 5F). We hypothesize that the resistosome-mediated Ca^2+^ influx releases the kinase domain of NbCDPKs from the calmodulin-like domain, resulting in NbRBOHB Ser-123 phosphorylation. Furthermore, nuclear-localized NbCDPKs may participate in the transcriptional upregulation of *NbRBOHB*. Our findings show the importance of the CDPK-RBOH module in the regulation of sustained ROS burst, which might be a connecting mechanism between effector recognition to downstream defense responses.

### Diverse functions and redundancy of CDPKs for RBOH activation

The phosphorylation of NbRBOHB Ser-123 was increased during ETI-like immune responses, which execute HR cell death (Fig. 3). Robust immune responses during ETI are accomplished by an increase in PTI signaling components transcriptionally and post-translationally (*4*). In Arabidopsis, RNA-seq analysis revealed that expression levels of several *CDPKs* are upregulated by chemical-induced expression of avirulence effector (*4*). Our results indicated that *NbCDPK4*, *NbCDPK5*, and *NbCDPK11* are transcriptionally upregulated in response to INF1 (Fig. 3H). WRKY-type transcription factors play pivotal roles in transcriptional reprogramming for plant immune responses (*40*). Indeed, *NbCDPK4* expression is upregulated by constitutively active mutants of MAPKK in a WRKY-dependent manner (*28*). In potato, changes in the expression level of 15 *CDPKs* were observed and the phosphorylation activities of CDPK are enhanced in response to pathogens (*41*). Given that overexpression of NbCDPK5 enhanced INF1-induced ROS production and the phosphorylation of NbRBOHB Ser-123 (Fig. 3F, G), transcriptional upregulation of *CDPKs* might be involved in the strong phosphorylation of the target site during ETI.

Multiple gene silencing of *NbCDPK4/5/9/10/11* attenuated NbRBOHB Ser-123 phosphorylation in response to INF1, but Ser-123 phosphorylation was still observed (Fig. 3D). Likewise, the phosphorylation of potato StRBOHB Ser-82 was also not affected by *NbCDPK4* and *NbCDPK5* knockdown in the heterologous expression system using *N. benthamiana* (*25*). Several studies identified the consensus motifs for CDPK-dependent phosphorylation (*42–44*). In addition to consensus motif sequence for substrate, subcellular localization of both substrates and CDPK are indispensable for the phosphorylation (*35*). To phosphorylate RBOH, CDPKs need to be anchored to the plasma membrane via myristoylation and S-acylation of the N-terminal V domain (*35*). Indeed, NbCDPK4 and NbCDPK5 containing myristoylation and S-acylation sites phosphorylated NbRBOHB, but NbCDPK9, NbCDPK10, and NbCDPK11 lacking myristoylation and S-acylation sites did not (Fig. 2D). Although myristoylation and S-acylation are necessary for plasma membrane localization of CDPK, acylated CDPKs do not always localize to the plasma membrane. For instance, the V domain of SlCDPK2 is responsible for the localization to the trans-Golgi network did not phosphorylate RBOH (*35*).

The CDPK gene family includes several paralogs that share a significant level of homology, which most likely emerged following a recent gene duplication event (*45*). The *N. benthamiana* genome encodes at least 80 CDPKs (Fig. 1A), of which about 50 contain myristoylation and S-acylation sites in their N-terminal V domain. Since Ser-123 phosphorylation was completely inhibited by the Ca^2+^-channel blocker, LaCl3 (Figs. 3D, 5D), suggesting that CDPKs are involved, but CDPKs that phosphorylate NbRBOHB could exist beside NbCDPK4 and NbCDPK5 identified in this study. In addition, NbRBOHB Ser-123 is shared with the consensus motif (LxRxxpS), which is phosphorylated by Snf1-related kinase (SnRK) (*46*). SnRK phosphorylates RBOH to enhance the tolerance to low nitrogen in tomato (*47*). We cannot rule out the possibility that diverse kinases, such as CDPKs, SnRK, RLCK, contribute to the regulation of RBOH activity cooperatively during ETI.

Multiple knockouts of *AtCPK4*, *AtCPK5*, *AtCPK6* and *AtCPK11* compromise flg22-induced PTI-ROS burst (*26*). The phosphorylation levels of AtRBOHD Ser-163, Ser-343, and Ser-347 are increased in response to flg22 and AvrRpt2 (*38*). AtRBOHD Ser-347 is shared with both CDPK- and RLCK-dependent phosphorylation motifs that are also conserved in NbRBOHB, implicating that phosphorylation of such motifs contributes to the ROS production during PTI and ETI. Our results indicated that NbRBOHB Ser-123 was not phosphorylated during PTI (Fig. 3C). The phosphorylation of AtRBOHD Ser-133, which corresponds to NbRBOHB Ser-123, during immune response, has not been determined by LC-MS/MS analysis due to the technical limitation (*18*). We think the possibility that the intensity of ROS production is regulated by the specific phosphorylation site dependent on signal inputs.

### CDPK-regulated ROS production and execution of HR cell death

The RBOH-dependent ROS production and spontaneous cell death are deeply connected. For instance, knockdown of *NbRBOHB* compromises HR cell death caused by INF1 and MEK2^DD^ transient expression (*16*). Transcription factor Alfin-like 7 positively regulates NLR-dependent HR cell death by suppressing the expression of ROS scavenging genes (*48*). CDPK might be involved in the induction of cell death through the regulation of RBOH activity. For example, Transgenic potato plants containing StCDPK5VK under the control of a pathogen-inducible promoter induce ROS accumulation and HR cell death in response to the virulent isolate of *P. infestans* (*27*). Moreover, ROS accumulation induced by the phosphorylation of OsRBOHH by OsCDPK5 and OsCDPK13 is required for programmed cell death to adopt a low oxygen environment in rice roots (*49*).

ROS are the second messenger to deliver the signal via oxidative post-translational modification (oxiPTM) of Cysteine (Cys) residues (*15*). NbRBOHB contributes to S-sulfenylation, one of the oxiPTM of cysteine, during plant immune response, which affects HR cell death (*50*). A recent study reported that S-sulfenylation of the transcription factor CCA1 HIKING EXPEDITION (CHE), in a RBOH-dependent manner, is required for the establishment of systemic acquired resistance (SAR) in local attack by avirulent strain of bacterial pathogen (*51*). These reports support that ROS can regulate immune responses including HR cell death by acting as intracellular- or intercellular second messengers. Furthermore, ROS lead to lipid peroxidation via iron-dependent Fenton reaction resulting in ferroptosis (*52*). A recent study showed that iron-, Ca^2+^- and ROS-dependent ferroptotic cell death is induced during rice immune responses (*53, 54*). CDPK-RBOH module might regulate HR cell death by oxiPTMs of proteins and execute ferroptosis with iron.

### CDPK is an important signaling hub connecting between Ca^2+^ and ROS

Upon PAMPs recognition, cyclic nucleotide-gated channels (CNGC) are activated by RLCK, and transient elevation of cytosolic Ca^2+^ is formed (*55*). Several other types of channels appear to contribute to the formation of the Ca^2+^ signature (*56*). Upon effector recognition, helper NLR forms a resistosome and causes Ca^2+^ influx at the plasma membrane (*11*). Recent progress indicates that following effector recognition by TNLs, their TIR-domain catalyzes the metabolism of NAD^+^, ATP, and other nucleic acids, and generates nucleotide metabolites acting as second messengers (*57*). These molecules induce the formation of helper NLR resistosome, ADR1 and NRG1, resulting in Ca^2+^ influx (*58*). Furthermore, mixed-lineage kinase domain-like (MLKL) contributes to Ca^2+^ influx downstream of TNLs activation (*59*). Thus, the elevation of cytosolic Ca^2+^ is a common event between PTI and ETI. MADA-motif conserved in the N-terminal of helper NLR is required for their Ca^2+^ channel activity and induction of HR cell death (*11, 13*). Our results showed that MADA-motif is also required for NbRBOHB-dependent ROS production and phosphorylation of NbRBOHB Ser-123 (Fig. 5). Notably, NbRBOHB Ser-337, which is predicted phosphorylation site by RLCK according to the phosphorylation motif SxxL (*60*), also phosphorylated in a Ca^2+^-dependent manner (Fig. 5B, D). These results indicate that CDPKs work as a hub connecting the Ca^2+^ channel activity of NLR and defense responses, such as ROS production. It is possible that CDPKs could be located upstream of RLCKs and initiate phosphorylation relay during ETI. Because sustained Ca^2+^ elevation is induced by any type of effector-NLR pair, the CDPK-RBOH module seems to commonly regulate ROS production and further downstream defense responses.

There are positive feedback circuits between ROS and Ca^2+^ (*61*). ROS signals are received by so called ROS sensors, such as receptor-like kinase, hydrogen-peroxide-induced Ca^2+^ increases (HPCA), and initiate Ca^2+^ elevation (*62*). On the other hand, ROS production by RBOH is activated by Ca^2+^ binding to the EF-hand motif in N-terminal (*23*). ROS production and Ca^2+^ influx induce each other, and as a result, cell-to-cell signal transduction, called ROS-Ca^2+^ wave, is formed (*61, 63*). This work showed that CDPK activated in a Ca^2+^-dependent manner, and phosphorylated RBOH to induce ROS production during ETI, supporting that the CDPK-RBOH module plays a pivotal role in the establishment of ROS-Ca^2+^ wave. AtCPK5 has been reported to contribute to the phosphorylation of AtRBOHD in the distal tissue (*64*). HR cell death is a local response at the attacked cells by avirulent pathogens, but ETI simultaneously induces systemic immune responses (*65*). Since RBOH plays a pivotal role in the establishment of SAR, the CDPK-RBOH module might be important to connect attacked cells with their surrounding cells, conferring effective defense responses.

## MATERIALS AND METHODS

### Plant materials and treatments

*N. benthamiana* plants were grown at 23°C under a 16-hour photoperiod and an 8-hour dark period in environmentally controlled growth cabinets.

### Agrobacterium-mediated transient expression in *N. benthamiana* (agroinfiltration)

The cDNAs of CDPKs were amplified by PCR from an *N. benthamiana* cDNA library as a template (*16*). The PCR products were cloned into pGEM-T Easy (Promega) and were sequenced. The CDPK and RBOH variants were generated using PrimeSTAR® Mutagenesis Basal Kit (TAKARA). CDPK and RBOH variants were cloned into pGreen binary vector (*66*) and pGD binary vector (*67*), respectively using the primers listed in the table S1. pGreen vector containing NtMEK2^DD^, pER8 vector carrying AVRblb2, and pICH47742 vector carrying NRC4 were described by Ishihama *et al.,* (*69*) and Adachi *et al.,* (*13, 28, 68*), respectively.

Agroinfiltration was done as described by Asai *et al.,* (*69*). These plasmids were transformed into Agrobacterium strain GV3101, which included the transformation helper plasmid pSoup (*70*) by electroporation. The overnight culture was diluted 3-fold in LB/kanamycin/rifampicin and was cultured for 2-3 hours. The cells were harvested by centrifugation and were resuspended in 10 mM MES-NaOH, pH 5.6, and 10 mM MgCl2. The suspensions were adjusted to OD600 = 0.5, and acetosyringone was added to 150 µM in final. The bacterial suspensions were incubated for 1 hour at room temperature and then, were infiltrated into leaves of 4-week-old *N. benthamiana* plants using a needleless syringe (*16*). Expression of AVRblb2 by estradiol-inducible promoter in pER8 was induced by 20 µM β-estradiol infiltration 24 hours after the agroinfiltration.

### Virus-induced gene silencing (VIGS)

Virus-induced gene silencing was done as described by Ratcliff *et al.* (*71*) and Ishihama *et al*. (*68*). The cDNA fragments were amplified from the *N. benthamiana* cDNA library using the primers listed in the table S1, and cloned into TRV vector pTV00 (*16*). The pTV00 vectors previously reported were used to silence NbRBOHB (*69*).

### Preparation of protein extracts

Microsomal proteins from *N. benthamiana* leaves were prepared as described by Kobayashi et al. with minor modifications (*27*). The leaves were thoroughly pulverized by Shake Master (Bio Medical Science), and mixed with protein extraction buffer (50 mM MOPS-KOH (pH 7.6), 0.5 M sorbitol, 10 mM DTT, 5 mM EGTA, 5 mM EDTA, 0.1 M NaF, 1 mM Na3VO4, 50 mM β-glycerophosphate, 0.1 mM AEBSF and 1 µM E-64), filtered through four layers of gauze, and centrifuged at 10,000*g* for 15 minutes at 4℃.

The supernatant was centrifuged at 150,000*g* for 40 minutes at 4℃. The pellet was suspended in suspension buffer (150 mM Tris-HCl (pH 7.5), 50 mM NaCl, 10% glycerol, 10 mM EDTA, 10 mM DTT, 1% IGEPAL CA630, 1 mM Na2MoO4, 2 mM NaF, 1 mM PMSF, 1.5 mM Na3VO4, 1% protease inhibitor cocktail for plant cell (Sigma Aldrich) and 50 mM β-glycerophosphate), and the suspension was used as microsomal proteins. The protein concentration was determined using the Protein Assay Dye Reagent (Bio-Rad) with BSA as a standard.

### Antibody production and immunoblotting

Immunoblotting was done using as primary antibodies against CDPKs-HA, NRC4s-6xHA (Clone H 9658; Sigma-Aldrich), FLAG-RBOH, FLAG-MEK2 (clone M2; Sigma-Aldrich), NbRBOHB pS123, pS337, and as secondary antibodies horseradish peroxidase-conjugated anti-rabbit or anti-mouse IgG antibody (Cytiva). The pS123 antibody and pS337 antibody for NbRBOHB were produced by Japan Bioserum Co., Ltd. using pS123 peptide (LVRNAS[PO3H2]SRIRQVSQELRRLA) and pS337 peptide (CSRNLS[PO3H2]QMLSQKLKH), respectively.

Equal amounts of proteins were separated on a 10% SDS-polyacrylamide gel and were transferred to a nitrocellulose membrane (Cytiva). After blocking in Block Ace (Yukizirushi) overnight at 4℃, the membranes were incubated with primary antibodies diluted with TBS-T (0.1 M Tris-HCl, pH 7.5, 0.15 M NaCl, 0.1% Tween 20) at room temperature for 1 hour. After washing with TBS-T, the membranes were incubated with horseradish peroxidase-conjugated secondary antibodies diluted with TBS-T for 1 hour at room temperature. The antibody-antigen complex was detected using the Super Signal West Dura Substrate (Thermo scientific) and the Light-Capture system (Luminograph III, ATTO), and immunostained bands were analyzed by the CS Analyzer 4 (ATTO). Protein loading control was stained using Ponceau 4R (*72*). The membranes were incubated in 5% acetic acid with 0.3% Ponceau 4R (Red food coloring; Kyoritsu-food Co., Ltd.), and washed in 1% acetic acid.

### RNA isolation and RT-PCR

Total RNA from *N. benthamiana* leaves was prepared using TRI Reagent® (Cosmo Bio Co., Ltd.) according to the manufacturer’s procedure. The reverse transcription was done using the ReverTra Ace qPCR RT kit (Toyobo). RT-qPCR was performed using the StepOnePlus Real-Time PCR system (Applied Biosystems) with THUNDERBIRD^TM^ SYBR^®^ qPCR Mix (TOYOBO). The expression of *NbRBOHB*, *NbWRKY8* and *NbCDPKs* were normalized to the expression of *EF-1α*. The gene-specific primers used for each sequence are listed in the table S1.

### Expression and purification of recombinant proteins

Unpurified proteins containing INF1 were prepared from *E. coli* carrying pFB53 (*inf1* in pFLAG-ATS) (*73*). The overnight culture of *E. coli* carrying pFB53 was diluted 50-fold in LB/ampicillin and was cultured to OD600 = 0.6. Then, 1 mM IPTG was added to the culture to induce *inf1* expression and it was further cultured for 4 hours. The culture was centrifuged, and its supernatant was filtrated and dialyzed in distilled water for 24 hours. Protein concentration was adjusted to 100 µg/mL.

pET30a (+) vectors containing NbRBOHB N-terminal fragments were transformed into *E. coli* cells of strain BL21-CodonPlus (DE3)-RIPL (Agilent). Expression of recombinant proteins were induced using Overnight Express^TM^ Autoinduction System1 (Novagen) according to the manufacturer’s procedure. Cells were harbored from overnight culture and resuspended in extraction buffer (20 mM HEPES-NaOH, pH 7.5).

The suspension was sonicated and centrifuged at 20,000*g* for 5 minutes at 4℃. The pellet was resuspended in extraction buffer containing 8 M urea and centrifuged at 20,000*g* for 5 minutes at 4℃. The supernatant was dialyzed against 20 mM HEPES-KOH, pH 7.6. pET32a (+) vectors containing NbCDPK5 was transformed into *E. coli* cells of strain BL21-Gold (DE3) pLysS (Stratagene). The overnight culture at 37℃ was transferred to 100-fold LB/ampicillin medium and cultured until OD600 was 0.6 at 37℃. Expression was induced by the addition of 0.4 mM IPTG for 3 hours at 28℃. Extraction and purification of CDPK proteins and Trx proteins were performed with Strep-Tactin Spin Columns (IBA).

### In vitro kinase assay

Kinase activity was determined in 15 µL of buffer (20 mM HEPES-KOH, pH 7.6, 1 mM DTT, and 5 mM MgCl2, 5 mM ATP with 0.1 mM CaCl2 or 2 mM EGTA) containing 0.7 mg of NbRBOHB N-terminal peptides and 70 ng of CDPKs incubated 30 minutes at room temperature. The reactions were stopped by the addition of SDS-PAGE sample buffer and incubation at 95℃ for 5 minutes.

### Measurement of ROS

Measurement of ROS in *N. benthamiana* leaves was performed as described by Kobayashi *et al*. (*25*). 0.5 mM L-012 in 10 mM MOPS-KOH, pH 7.4, was infiltrated into *N. benthamiana* leaves using a needleless syringe. The chemiluminescence was monitored continuously by a Light-Capture system (Luminograph III, ATTO) for 5 minutes. The intensity of chemiluminescence was quantified using CS Analyzer 4 (ATTO).

## Supplementary materials

**Fig. S1.**
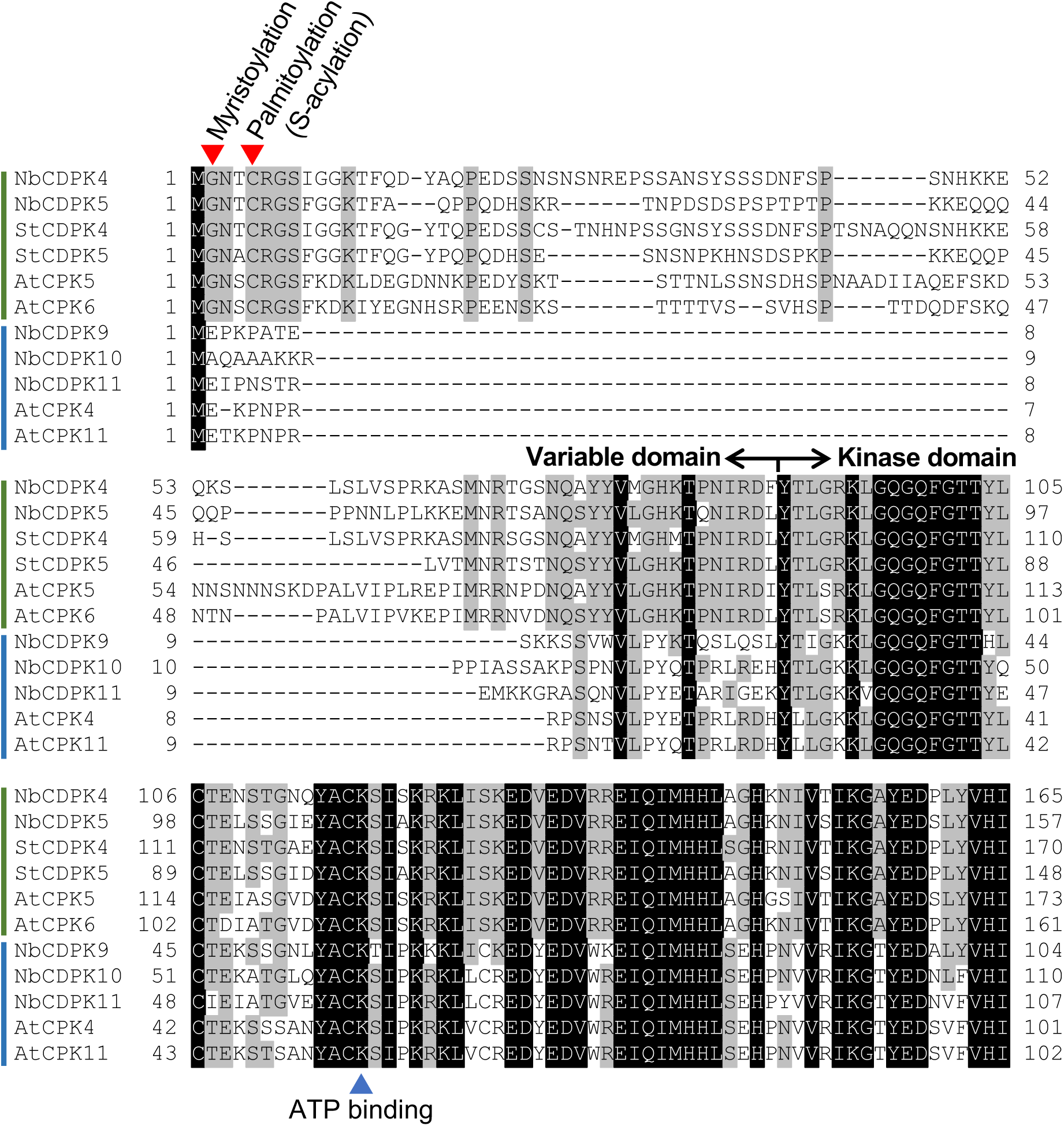
Multiple alignments of the N-terminal region of CDPKs. The amino acid sequences of N-terminal regions of CDPKs, Arabiodopsis AtCPK4, AtCPK5, AtCPK6, AtCPK11, potato StCDPK4, StCDPK5, and Nicotiana benthamiana NbCDPK4, NbCDPK5, NbCDPK9, NbCDPK10, NbCDPK11 were aligned by ClustalW. Red arrowheads indicate the predicted myristoylation and S-acylation sites. Blue arrowheads indicate the ATP binding sites.

**Fig. S2.**
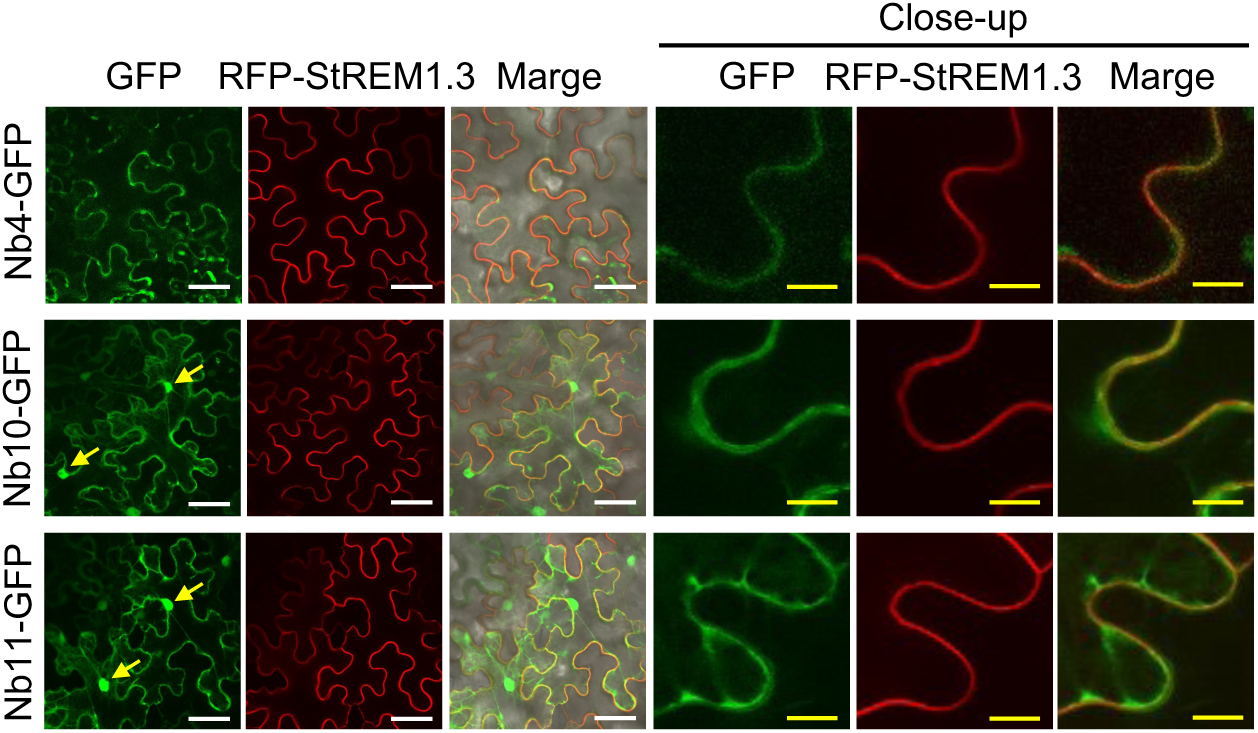
Subcellular localization of NbCDPK4, NbCDPK10, and NbCDPK11. GFP-fused NbCDPKs were transiently co-expressed with RFP-StREM1.3 via agroinfiltration. GFP- and RFP-fluorescence were observed under confocal microscope 3 days after agroinfiltration. Arrows indicate the nucleus. White bars = 50 μm, yellow bars = 10 μm.

**Fig S3.**
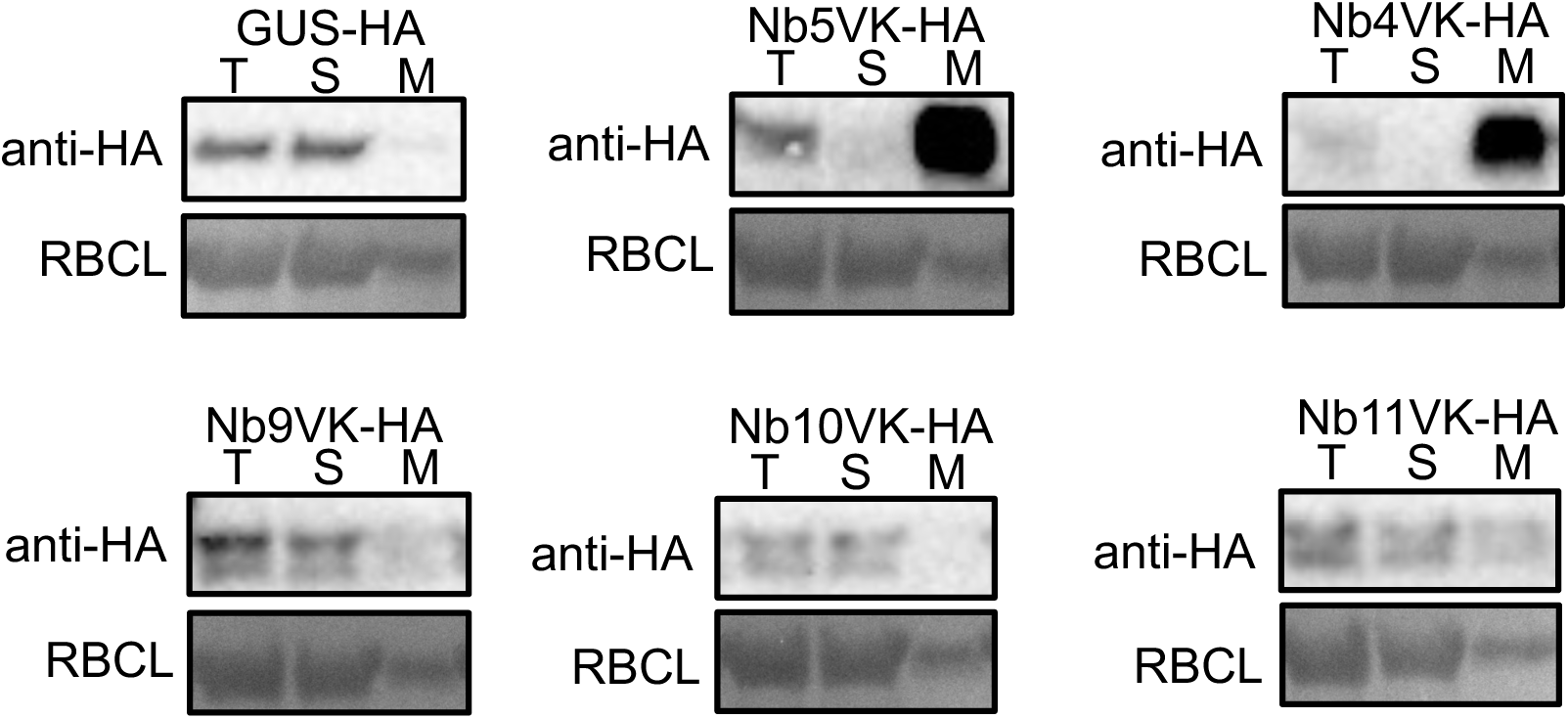
NbCDPK4 and NbCDPK5 were enriched in microsome fraction. *A. tumefaciens* harboring HA-tagged NbCDPKVKs were infiltrated into *N. benthamiana* leaves. Two days after agroinfiltration, total protein (T), supernatant fraction (S), and microsome fraction (M) were prepared. Immunoblot analyses were performed using anti-HA antibody. Protein loads were monitored by MemCode staining of the bands corresponding to Rubisco large subunit (RBCL).

**Fig. S4.**
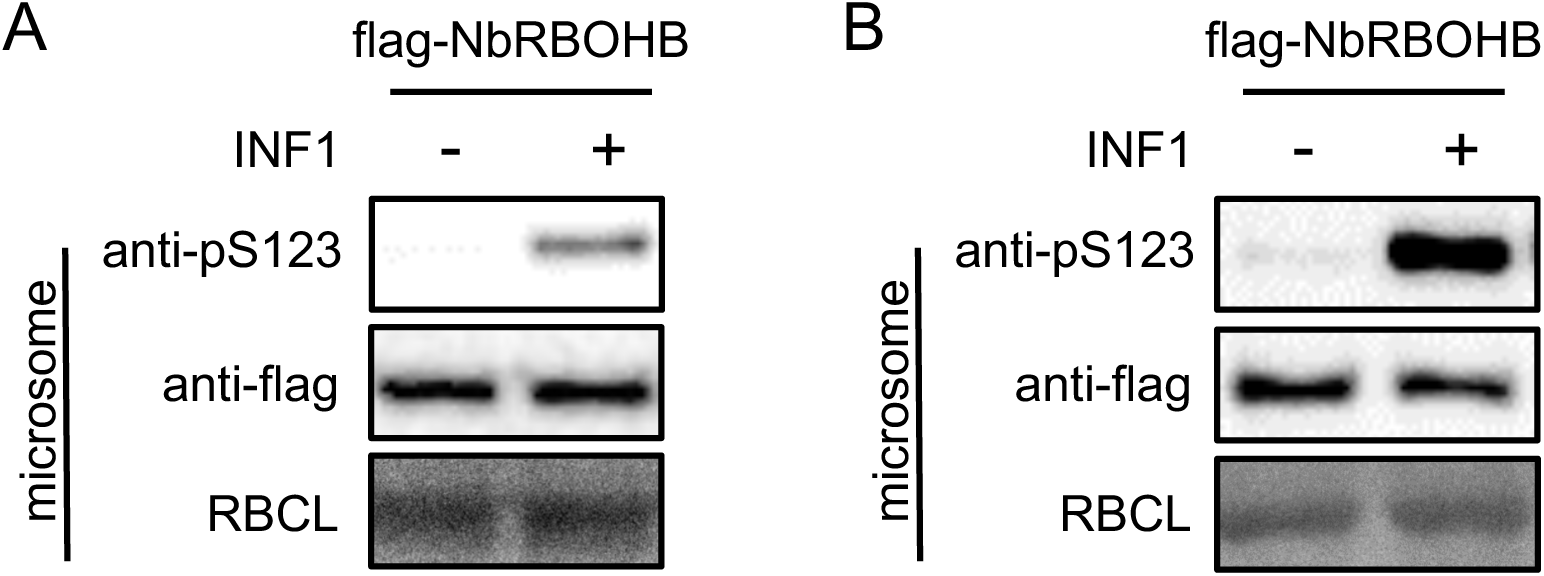
The phosphorylation of NbRBOHB Ser-123 induced by INF1. (A and B) Phosphorylation of NbRBOHB Ser-123 in response to INF1 peptide infiltration. NbRBOHB were transiently expressed by agroinfiltration. Two days after agroinfiltration, the leaves were treated with INF1 peptide. Microsomal fractions were prepared (A) 12 hours and (B) 24 hours after INF1 treatment. Immunoblot analyses were performed using anti-pS123 and anti-flag antibodies. Protein loads were monitored by ponceau 4R staining of the bands corresponding to RBCL.

**Fig. S5.**
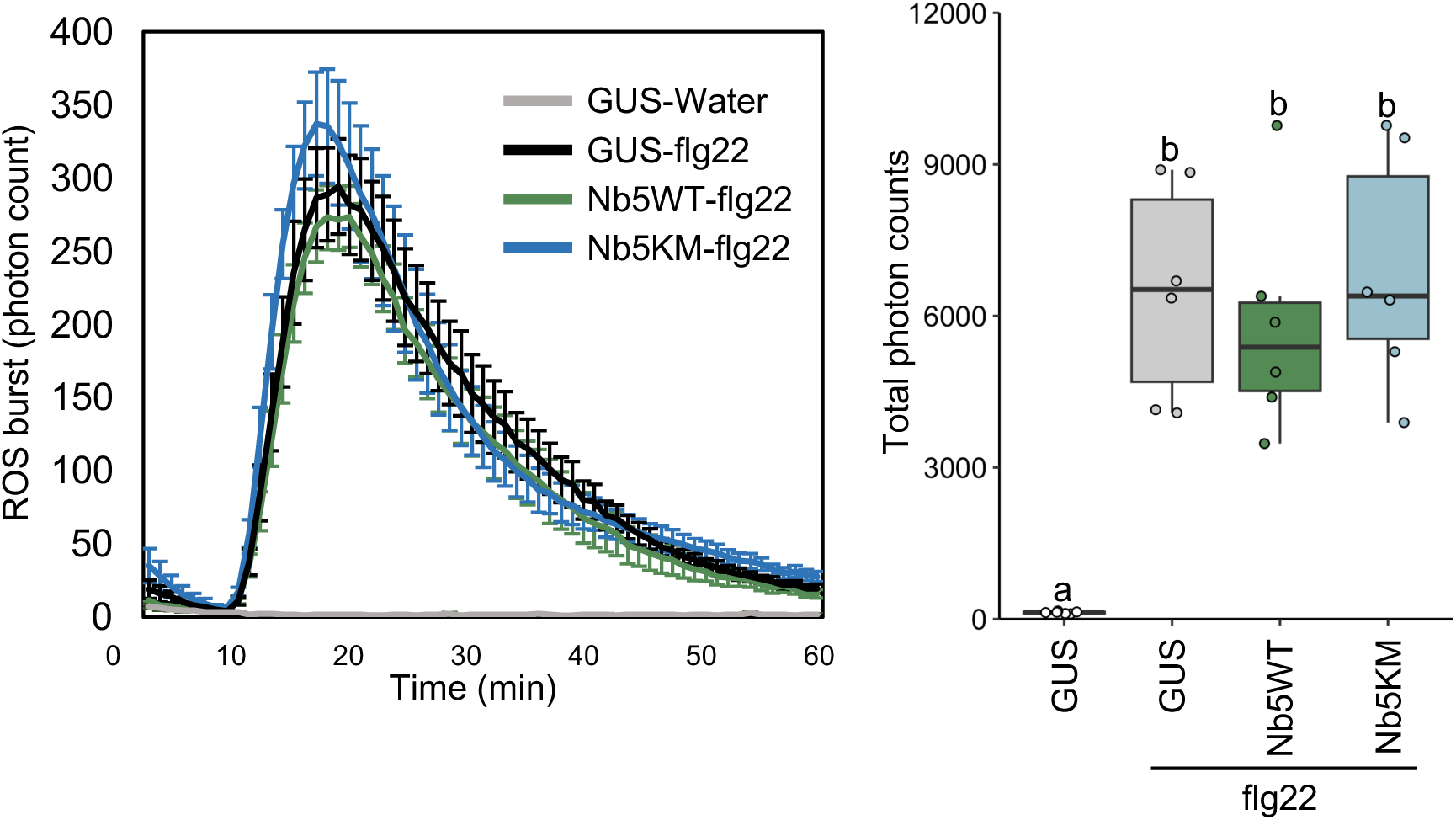
Overexpression of NbCDPK5 did not affect PTI-ROS burst. flg22-triggered ROS bursts in *N. benthamiana* leaves overexpressing NbCDPK5 variants or GUS control two days after agroinfiltration. ROS production was measured by monitoring chemiluminescence of luminol every minute for an hour. Letters represent each significance group, determined through Tukey’s multiple range test.

**Fig. S6.**
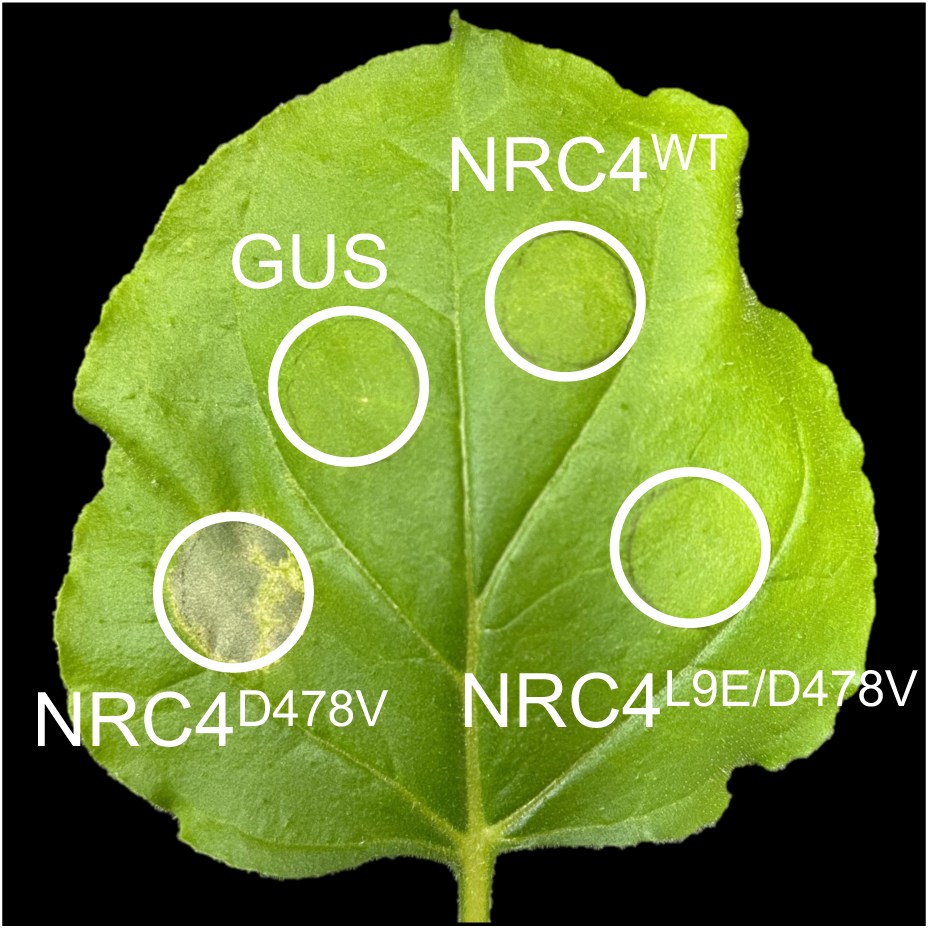
HR cell death induced by transient expression of NRC4 variants. L9E single mutation (NRC4^D478V/L9E^) impaired cell death by NRC4^DV^. NRC4^WT^-6xHA, NRC4^D478V^-6xHA, NRC4^D478V/L9E^-6xHA and GUS-HA were expressed and photographed at 5 days after agroinfiltration.

---

**Supplemental table 1.**
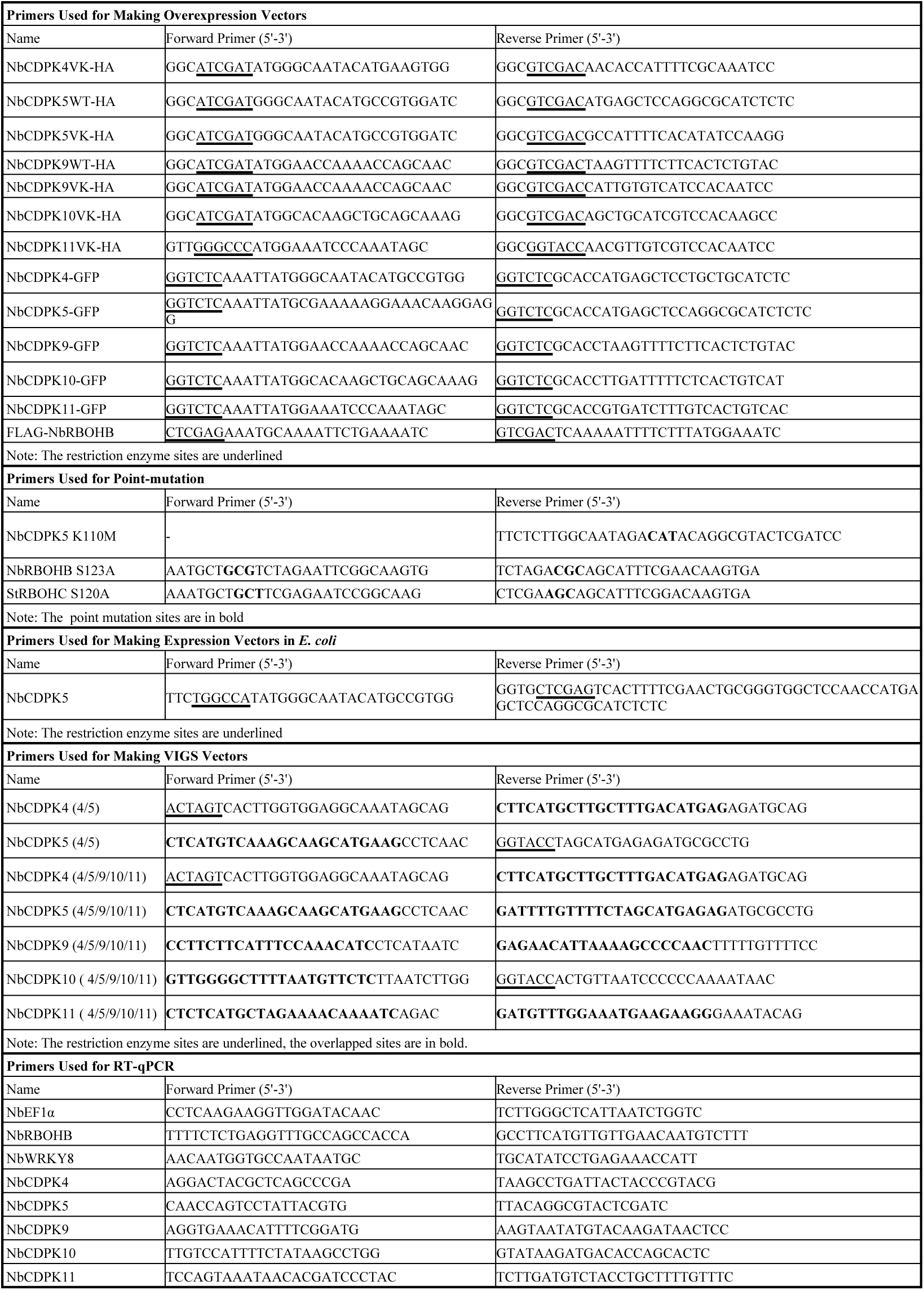
Primers used for this study.

## Acknowledgments

We thank Phil Mullineaux and Roger Hellens for pGreen vector, Andrew O. Jackson for pGD binary vector, Sophien Kamoun for INF1 and AVRblb2/Rpiblb2 constructs, David C. Baulcombe for TRV vector and p19 construct, and the Leaf Tobacco Research Center, Japan, for *N. benthamiana* seeds. We also thank Mitsuhiro Yada and Yutaro Shiraishi for technical assistance with cloning for NbCDPK and constructing expression vectors.

## Author contributions

Y.H. performed measurement of ROS, immunoblotting for detecting phosphorylation of NbRBOHB in plants, RT-qPCR, co-IP assay, constructing expression vectors, and analysis of the subcellular localization. M.Y. and M.Y. performed cloning for CDPK and RBOH, constructing expression vectors, and *in vitro* kinase assay. H.A and H.Y. designed experiments and supervised research. Y.H and H.Y wrote the manuscript.

## Data availability

All data are available from the corresponding author upon requests.

## References

1. J. D. Jones, J. L. Dangl, The plant immune system. Nature 444, 323–329 (2006).

2. S. Lacombe, A. Rougon-Cardoso, E. Sherwood, N. Peeters, D. Dahlbeck, H. P. van Esse, M. Smoker, G. Rallapalli, B. P. Thomma, B. Staskawicz, J. D. Jones, C. Zipfel, Interfamily transfer of a plant pattern-recognition receptor confers broad-spectrum bacterial resistance. Nat Biotechnol 28, 365–369 (2010).

3. M. Yuan, Z. Jiang, G. Bi, K. Nomura, M. Liu, Y. Wang, B. Cai, J. M. Zhou, S. Y. He, X. F. Xin, Pattern-recognition receptors are required for NLR-mediated plant immunity. Nature 592, 105–109 (2021).

4. B. P. M. Ngou, H. K. Ahn, P. Ding, J. D. G. Jones, Mutual potentiation of plant immunity by cell-surface and intracellular receptors. Nature 592, 110–115 (2021).

5. X. Q. Yu, H. Q. Niu, C. Liu, H. L. Wang, W. Yin, X. Xia, PTI-ETI synergistic signal mechanisms in plant immunity. Plant Biotechnol J 22, 2113–2128 (2024).

6. F. Locci, J. E. Parker, Plant NLR immunity activation and execution: a biochemical perspective. Open Biol 14, 230387 (2024).

7. S. Huang, E. Li, F. Jia, Z. Han, J. Chai, Assembly and functional mechanisms of plant NLR resistosomes. Curr Opin Struct Biol 90, 102977 (2025).

8. G. Bi, M. Su, N. Li, Y. Liang, S. Dang, J. Xu, M. Hu, J. Wang, M. Zou, Y. Deng, Q. Li, S. Huang, J. Li, J. Chai, K. He, Y. H. Chen, J. M. Zhou, The ZAR1 resistosome is a calcium-permeable channel triggering plant immune signaling. Cell 184, 3528–3541.e3512 (2021).

9. H. Adachi, L. Derevnina, S. Kamoun, NLR singletons, pairs, and networks: evolution, assembly, and regulation of the intracellular immunoreceptor circuitry of plants. Curr Opin Plant Biol 50, 121–131 (2019).

10. C. H. Wu, A. Abd-El-Haliem, T. O. Bozkurt, K. Belhaj, R. Terauchi, J. H. Vossen, S. Kamoun, NLR network mediates immunity to diverse plant pathogens. Proc Natl Acad Sci U S A 114, 8113–8118 (2017).

11. F. Liu, Z. Yang, C. Wang, Z. You, R. Martin, W. Qiao, J. Huang, P. Jacob, J. L. Dangl, J. E. Carette, S. Luan, E. Nogales, B. J. Staskawicz, Activation of the helper NRC4 immune receptor forms a hexameric resistosome. Cell 187, 4877–4889.e4815 (2024).

12. J. Wang, M. Hu, J. Qi, Z. Han, G. Wang, Y. Qi, H. W. Wang, J. M. Zhou, J. Chai, Reconstitution and structure of a plant NLR resistosome conferring immunity. Science 364, (2019).

13. H. Adachi, M. P. Contreras, A. Harant, C. H. Wu, L. Derevnina, T. Sakai, C. Duggan, E. Moratto, T. O. Bozkurt, A. Maqbool, J. Win, S. Kamoun, An N-terminal motif in NLR immune receptors is functionally conserved across distantly related plant species. Elife 8, (2019).

14. R. Mittler, ROS are good. Trends Plant Sci 22, 11–19 (2017).

15. S. Kimura, C. Waszczak, K. Hunter, M. Wrzaczek, Bound by fate: The role of reactive oxygen species in receptor-like kinase signaling. Plant Cell 29, 638–654 (2017).

16. H. Yoshioka, N. Numata, K. Nakajima, S. Katou, K. Kawakita, O. Rowland, J. D. Jones, N. Doke, *Nicotiana benthamiana* gp91*^phox^* homologs *NbrbohA* and *NbrbohB* participate in H2O2 accumulation and resistance to *Phytophthora infestans*. Plant Cell 15, 706–718 (2003).

17. B. Castro, M. Citterico, S. Kimura, D. M. Stevens, M. Wrzaczek, G. Coaker, Stress-induced reactive oxygen species compartmentalization, perception and signalling. Nat Plants 7, 403–412 (2021).

18. Y. Kadota, J. Sklenar, P. Derbyshire, L. Stransfeld, S. Asai, V. Ntoukakis, J. D. Jones, K. Shirasu, F. Menke, A. Jones, C. Zipfel, Direct regulation of the NADPH oxidase RBOHD by the PRR-associated kinase BIK1 during plant immunity. Mol Cell 54, 43–55 (2014).

19. S. Kimura, K. Hunter, L. Vaahtera, H. C. Tran, M. Citterico, A. Vaattovaara, A. Rokka, S. C. Stolze, A. Harzen, L. Meißner, M. M. T. Wilkens, T. Hamann, M. Toyota, H. Nakagami, M. Wrzaczek, CRK2 and C-terminal phosphorylation of NADPH oxidase RBOHD regulate reactive oxygen species production in Arabidopsis. Plant Cell 32, 1063–1080 (2020).

20. L. Kong, X. Ma, C. Zhang, S. I. Kim, B. Li, Y. Xie, I. C. Yeo, H. Thapa, S. Chen, T. P. Devarenne, T. Munnik, P. He, L. Shan, Dual phosphorylation of DGK5-mediated PA burst regulates ROS in plant immunity. Cell 187, 609–623.e621 (2024).

21. F. Qi, J. Li, Y. Ai, K. Shangguan, P. Li, F. Lin, Y. Liang, DGK5β-derived phosphatidic acid regulates ROS production in plant immunity by stabilizing NADPH oxidase. Cell Host Microbe 32, 425–440.e427 (2024).

22. C. Zhang, K. E. Atanasov, R. Alcázar, Spermine inhibits PAMP-induced ROS and Ca^2+^ burst and reshapes the transcriptional landscape of PAMP-triggered immunity in Arabidopsis. J Exp Bot 74, 427–442 (2023).

23. M. Sagi, R. Fluhr, Superoxide production by plant homologues of the gp91*^phox^* NADPH oxidase. Modulation of activity by calcium and by tobacco mosaic virus infection. Plant Physiol 126, 1281–1290 (2001).

24. T. Yip Delormel, M. Boudsocq, Properties and functions of calcium-dependent protein kinases and their relatives in *Arabidopsis thaliana*. New Phytol 224, 585–604 (2019).

25. M. Kobayashi, I. Ohura, K. Kawakita, N. Yokota, M. Fujiwara, K. Shimamoto, N. Doke, H. Yoshioka, Calcium-dependent protein kinases regulate the production of reactive oxygen species by potato NADPH oxidase. Plant Cell 19, 1065–1080 (2007).

26. M. Boudsocq, M. R. Willmann, M. McCormack, H. Lee, L. Shan, P. He, J. Bush, S. H. Cheng, J. Sheen, Differential innate immune signalling via Ca^2+^ sensor protein kinases. Nature 464, 418–422 (2010).

27. M. Kobayashi, M. Yoshioka, S. Asai, H. Nomura, K. Kuchimura, H. Mori, N. Doke, H. Yoshioka, StCDPK5 confers resistance to late blight pathogen but increases susceptibility to early blight pathogen in potato via reactive oxygen species burst. New Phytol 196, 223–237 (2012).

28. H. Adachi, T. Nakano, N. Miyagawa, N. Ishihama, M. Yoshioka, Y. Katou, T. Yaeno, K. Shirasu, H. Yoshioka, WRKY transcription factors phosphorylated by MAPK regulate a plant immune NADPH oxidase in *Nicotiana benthamiana*. Plant Cell 27, 2645–2663 (2015).

29. M. Boudsocq, J. Sheen, CDPKs in immune and stress signaling. Trends Plant Sci 18, 30–40 (2013).

30. J. F. Harper, G. Breton, A. Harmon, Decoding Ca^2+^ signals through plant protein kinases. Annu Rev Plant Biol 55, 263–288 (2004).

31. J. Harper, J. Huang, S. Lloyd, Genetic indentification of an autoinhibitor in CDPK, a protein-kinase with a calmodulin-like domain. Biochemistry 33, 7267–7277 (1994).

32. A. Ludwig, H. Saitoh, G. Felix, G. Freymark, O. Miersch, C. Wasternack, T. Boller, J. Jones, T. Romeis, Ethylene-mediated cross-talk between calcium-dependent protein kinase and MAPK signaling controls stress responses in plants. Proc Natl Acad Sci U S A 102, 10736–10741 (2005).

33. M. Kapiloff, J. Mathis, C. Nelson, C. Lin, M. Rosenfeld, Calcium calmodulin-dependent protein-kinase mediates a pathway for transcriptional regulation. Proc Natl Acad Sci U S A 88, 3710–3714 (1991).

34. J. Sheen, Ca^2+^-dependent protein kinases and stress signal transduction in plants. Science 274, 1900–1902 (1996).

35. S. Asai, T. Ichikawa, H. Nomura, M. Kobayashi, Y. Kamiyoshihara, H. Mori, Y. Kadota, C. Zipfel, J. D. G. Jones, H. Yoshioka, The variable domain of a plant calcium-dependent protein kinase (CDPK) confers subcellular localization and substrate recognition for NADPH oxidase. J Biol Chem 288, 14332–14340 (2013).

36. A. Perraki, J. Cacas, J. Crowet, L. Lins, M. Castroviejo, S. German-Retana, S. Mongrand, S. Raffaele, Plasma membrane localization of *Solanum tuberosum* remorin from group 1, homolog 3 is mediated by conformational changes in a novel C-terminal anchor and required for the restriction of potato virus X movement. Plant Physiol 160, 624–637 (2012).

37. S. Imano, M. Fushimi, M. Camagna, A. Tsuyama-Koike, H. Mori, A. Ashida, A. Tanaka, I. Sato, S. Chiba, K. Kawakita, M. Ojika, D. Takemoto, AP2/ERF transcription factor NbERF-IX-33 is involved in the regulation of phytoalexin production for the resistance of *Nicotiana benthamiana* to *Phytophthora infestans*. Front Plant Sci 12, 821574 (2021).

38. Y. Kadota, T. W. H. Liebrand, Y. Goto, J. Sklenar, P. Derbyshire, F. L. H. Menke, M. A. Torres, A. Molina, C. Zipfel, G. Coaker, K. Shirasu, Quantitative phosphoproteomic analysis reveals common regulatory mechanisms between effector- and PAMP-triggered immunity in plants. New Phytol 221, 2160–2175 (2019).

39. R. Takabatake, Y. Ando, S. Seo, S. Katou, S. Tsuda, Y. Ohashi, I. Mitsuhara, MAP kinases function downstream of HSP90 and upstream of mitochondria in TMV resistance gene N-mediated hypersensitive cell death. Plant Cell Physiol 48, 498–510 (2007).

40. T. Javed, S. J. Gao, WRKY transcription factors in plant defense. Trends Genet 39, 787–801 (2023).

41. R. Gromadka, J. Ciesla, K. Olszak, J. Szczegielniak, G. Muszynska, L. Polkowska-Kowalczyk, Genome-wide analysis and expression profiling of calcium-dependent protein kinases in potato (*Solanum tuberosum*). Plant Growth Regul 84, 303–315 (2018).

42. J. Huang, S. Hardin, S. Huber, Identification of a novel phosphorylation motif for CDPKs: Phosphorylation of synthetic peptides lacking basic residues at P-3/P-4. Arch Biochem Biophys 393, 61–66 (2001).

43. J. Huang, S. Huber, Phosphorylation of synthetic peptides by a CDPK and plant SNF1-related protein kinase. Influence of proline and basic amino acid residues at selected positions. Plant Cell Physiol 42, 1079–1087 (2001).

44. A. Curran, I. Chang, C. Chang, S. Garg, R. Miguel, Y. Barron, Y. Li, S. Romanowsky, J. Cushman, M. Gribskov, A. Harmon, J. Harper, Calcium-dependent protein kinases from *Arabidopsis* show substrate specificity differences in an analysis of 103 substrates. Front Plant Sci 2, (2011).

45. T. Mohanta, N. Mohanta, Y. Mohanta, H. Bae, Genome-wide identification of calcium dependent protein kinase gene family in plant lineage shows presence of novel D-x-D and D-E-L motifs in EF-hand domain. Front Plant Sci 6, (2015).

46. F. Vlad, B. E. Turk, P. Peynot, J. Leung, S. Merlot, A versatile strategy to define the phosphorylation preferences of plant protein kinases and screen for putative substrates. Plant J 55, 104–117 (2008).

47. X. Zheng, H. Yang, J. Zou, W. Jin, Z. Qi, P. Yang, J. Yu, J. Zhou, SnRK1α1-mediated RBOH1 phosphorylation regulates reactive oxygen species to enhance tolerance to low nitrogen in tomato. Plant Cell 37, (2024).

48. D. Zhang, Z. Gao, H. Zhang, Y. Yang, X. Yang, X. Zhao, H. Guo, U. Nagalakshmi, D. Li, S. P. Dinesh-Kumar, Y. Zhang, The MAPK-Alfin-like 7 module negatively regulates ROS scavenging genes to promote NLR-mediated immunity. Proc Natl Acad Sci U S A 120, e2214750120 (2023).

49. J. Li, T. Ishii, M. Yoshioka, Y. Hino, M. Nomoto, Y. Tada, H. Yoshioka, H. Takahashi, T. Yamauchi, M. Nakazono, CDPK5 and CDPK13 play key roles in acclimation to low oxygen through the control of RBOH-mediated ROS production in rice. Plant Physiol 197, (2024).

50. Y. Hino, T. Inada, M. Yoshioka, H. Yoshioka, NADPH oxidase-mediated sulfenylation of cysteine derivatives regulates plant immunity. J Exp Bot 75, 4641–4654 (2024).

51. L. Cao, S. Karapetyan, H. Yoo, T. Chen, M. Mwimba, X. Zhang, X. Dong, H2O2 sulfenylates CHE, linking local infection to the establishment of systemic acquired resistance. Science 385, 1211–1217 (2024).

52. S. Dixon, K. Lemberg, M. Lamprecht, R. Skouta, E. Zaitsev, C. Gleason, D. Patel, A. Bauer, A. Cantley, W. Yang, B. Morrison, B. Stockwell, Ferroptosis: an iron-dependent form of nonapoptotic cell death. Cell 149, 1060–1072 (2012).

53. J. Wang, W. G. Choi, N. K. Nguyen, D. Liu, S. H. Kim, D. Lim, B. K. Hwang, N. S. Jwa, Cytoplasmic Ca^2+^ influx mediates iron- and reactive oxygen species-dependent ferroptotic cell death in rice immunity. Front Plant Sci 15, 1339559 (2024).

54. Q. Shen, M. Liang, F. Yang, Y. Deng, N. Naqvi, Ferroptosis contributes to developmental cell death in rice blast. New Phytol 227, 1831–1846 (2020).

55. W. Tian, C. Hou, Z. Ren, C. Wang, F. Zhao, D. Dahlbeck, S. Hu, L. Zhang, Q. Niu, L. Li, B. J. Staskawicz, S. Luan, A calmodulin-gated calcium channel links pathogen patterns to plant immunity. Nature 572, 131–135 (2019).

56. P. Köster, T. A. DeFalco, C. Zipfel, Ca^2+^ signals in plant immunity. EMBO J 41, e110741 (2022).

57. A. Jia, S. Huang, S. Ma, X. Chang, Z. Han, J. Chai, TIR-catalyzed nucleotide signaling molecules in plant defense. Curr Opin Plant Biol 73, 102334 (2023).

58. P. Jacob, N. Kim, F. Wu, F. El Kasmr, Y. Chi, W. Walton, O. Furzer, A. Lietzan, S. Sunil, K. Kempthorn, M. Redinbo, Z. Pei, L. Wan, J. Dangl, Plant “helper” immune receptors are Ca^2+^-permeable nonselective cation channels. Science 373, 420-+ (2021).

59. Q. Shen, K. Hasegawa, N. Oelerich, A. Prakken, L. W. Tersch, J. Wang, F. Reichhardt, A. Tersch, J. C. Choo, T. Timmers, K. Hofmann, J. E. Parker, J. Chai, T. Maekawa, Cytoplasmic calcium influx mediated by plant MLKLs confers TNL-triggered immunity. Cell Host Microbe 32, 453–465.e456 (2024).

60. J. Chu, I. Monte, T. A. DeFalco, P. Köster, P. Derbyshire, F. L. H. Menke, C. Zipfel, Conservation of the PBL-RBOH immune module in land plants. Curr Biol 33, 1130–1137.e1135 (2023).

61. S. Gilroy, N. Suzuki, G. Miller, W. G. Choi, M. Toyota, A. R. Devireddy, R. Mittler, A tidal wave of signals: calcium and ROS at the forefront of rapid systemic signaling. Trends Plant Sci 19, 623–630 (2014).

62. F. Wu, Y. Chi, Z. Jiang, Y. Xu, L. Xie, F. Huang, D. Wan, J. Ni, F. Yuan, X. Wu, Y. Zhang, L. Wang, R. Ye, B. Byeon, W. Wang, S. Zhang, M. Sima, S. Chen, M. Zhu, J. Pei, D. M. Johnson, S. Zhu, X. Cao, C. Pei, Z. Zai, Y. Liu, T. Liu, G. B. Swift, W. Zhang, M. Yu, Z. Hu, J. N. Siedow, X. Chen, Z. M. Pei, Hydrogen peroxide sensor HPCA1 is an LRR receptor kinase in Arabidopsis. Nature 578, 577–581 (2020).

63. Y. Fichman, R. Mittler, Integration of electric, calcium, reactive oxygen species and hydraulic signals during rapid systemic signaling in plants. Plant J 107, 7–20 (2021).

64. U. Dubiella, H. Seybold, G. Durian, E. Komander, R. Lassig, C. P. Witte, W. X. Schulze, T. Romeis, Calcium-dependent protein kinase/NADPH oxidase activation circuit is required for rapid defense signal propagation. Proc Natl Acad Sci U S A 110, 8744–8749 (2013).

65. J. A. Nie, X. H. Ding, X. R. Zhong, W. C. Shi, Z. Gao, Transcellular regulation of ETI-induced cell death. Trends Plant Sci, (2025).

66. R. Hellens, E. Edwards, N. Leyland, S. Bean, P. Mullineaux, pGreen:: a versatile and flexible binary Ti vector for *Agrobacterium*-mediated plant transformation. Plant Mol Biol 42, 819–832 (2000).

67. M. Goodin, R. Dietzgen, D. Schichnes, S. Ruzin, A. Jackson, pGD vectors: versatile tools for the expression of green and red fluorescent protein fusions in agroinfiltrated plant leaves. Plant J 31, 375–383 (2002).

68. N. Ishihama, R. Yamada, M. Yoshioka, S. Katou, H. Yoshioka, Phosphorylation of the *Nicotiana benthamiana* WRKY8 transcription factor by MAPK functions in the defense response. Plant Cell 23, 1153–1170 (2011).

69. S. Asai, K. Ohta, H. Yoshioka, MAPK signaling regulates nitric oxide and NADPH oxidase-dependent oxidative bursts in *Nicotiana benthamiana*. Plant Cell 20, 1390–1406 (2008).

70. R. Hellens, P. Mullineaux, H. Klee, Technical Focus:a guide to Agrobacterium binary Ti vectors. Trends Plant Sci 5, 446–451 (2000).

71. F. Ratcliff, A. M. Martin-Hernandez, D. C. Baulcombe, Technical advance. Tobacco rattle virus as a vector for analysis of gene function by silencing. Plant J 25, 237–245 (2001).

72. M. Contreras, H. Pai, R. Thompson, C. Marchal, J. Claeys, H. Adachi, S. Kamoun, The nucleotide-binding domain of NRC-dependent disease resistance proteins is sufficient to activate downstream helper NLR oligomerization and immune signaling. New Phytol 243, 345–361 (2024).

73. S. Kamoun, P. van West, A. J. de Jong, K. E. de Groot, V. G. Vleeshouwers, F. Govers, A gene encoding a protein elicitor of *Phytophthora infestans* is down-regulated during infection of potato. Mol Plant Microbe Interact 10, 13–20 (1997).

